# Sensory integration, temporal prediction and rule discovery reflect interdependent inference processes

**DOI:** 10.1101/2025.04.29.651167

**Authors:** Lucas Benjamin, Benjamin Morillon, Valentin Wyart

## Abstract

Deciphering the structure of variable sensory input is key to building an accurate model of one’s environment. Humans can integrate stimulus sequences to estimate their sensory statistics, predict the timing of upcoming stimuli, but also discover rules governing sequence generation. However, whether these three forms of inference operate independently or synergistically remains untested. Here we report selective interactions between sensory integration, temporal prediction, and rule discovery in humans. Participants were exposed to rhythmic sequences of 10 stimuli governed or not by a latent rule – a predictable change in stimulus statistics after 5 stimuli – and then asked to predict the 10^th^ stimulus from incomplete sequences. Individual differences in sensory integration timescale for rule-free sequences predicted efficient rule discovery. Conversely, discovering the latent rule shaped the timescale and format of sensory integration for rule-based sequences. Tampering with the rhythmicity of stimulus presentation impaired rule discovery without affecting sensory integration accuracy. Selective perturbations of recurrent neural networks (RNNs) trained in the same conditions confirmed these specific interactions. Together, these findings provide novel insights into the flexibility of human inferences based on variable yet predictable sensory input.

## Introduction

The sensory environments we live in are variable, yet highly organized. Understanding the generative organization that governs their structure is crucial for reducing uncertainty. To do so, humans rely on multiple inference abilities including sensory integration^1–4^, temporal prediction^5–8^, categorization based on arbitrary mappings of sensory features^9–11^, learning statistical dependencies between stimuli^12–18^, and the learning of hidden rules describing the patterns of relations between successive stimuli^19–23^. In a majority of studies, these different learning abilities are assumed, either implicitly or explicitly, to rely on different cognitive and neural mechanisms. But some studies, on the opposite, suggest a single unified cognitive apparatus for those different learning processes^24^. This debate lacks empirical data, and these abilities themselves have rarely been studied conjointly, resulting in a fragmented description of human learning abilities. Here, we studied three humans inference capacities conjointly and empirically tested if they rely on similar or distinct and independent or interrelated cognitive processes: sensory integration, temporal prediction, and rule discovery.

So far, inference capacities have mostly been studied in isolation, targeting different research questions and using different methods. In perceptual decisions research, studies have examined how sensory information is integrated across sequences of stimuli – or across samples from the same stimulus – to reach a decision^3,4,25,26^. We refer to this ability as *sensory integration*. This process is known to be prone to a number of suboptimalities constraining the accuracy of perceptual decisions across individuals^27–30^. Yet, it remains largely unknown whether these sensory integration suboptimalities are confined to sensory integration itself, or whether they act as a broader cognitive bottleneck impairing higher-order inferences, such as rule discovery.

To investigate this possible effect of sensory integration limitations on higher-order inferences, we designed a rule-based visual prediction task that requires both sensory integration and the discovery of a hidden rule for accurate predictions. Rule discovery is known to depend on the rule’s algorithmic complexity and to vary across individuals^22,31^. However, except for a few specific studies^32^, sequences used in most rule discovery tasks are typically stripped of any variability, eliminating the need to integrate sensory information across stimuli. In our rule-based prediction task, we used rhythmic isochronous sequences of 10 oriented stimuli (**Fig. 1A**) sampled from probability distributions whose mean orientations switch (i.e., rotate by 90 degrees) after 5 stimuli – a latent rule that was not described to participants. After exposing participants to full sequences, we asked them to predict the 10^th^ stimulus from incomplete sequences of 3, 5, 7 or 9 stimuli (*switch* blocks, **Fig. 1B**). In other blocks (*static* blocks), we used sequences where all stimuli were drawn from a single distribution – without a latent rule – to investigate if suboptimalities in sensory integration measured in static blocks could impact discovery of the latent rule in switch blocks. By introducing inter-stimulus temporal jitter, we further investigated whether a third inference ability – temporal prediction – affected either of these two forms of inference, or their interrelation.

**Figure 1.**
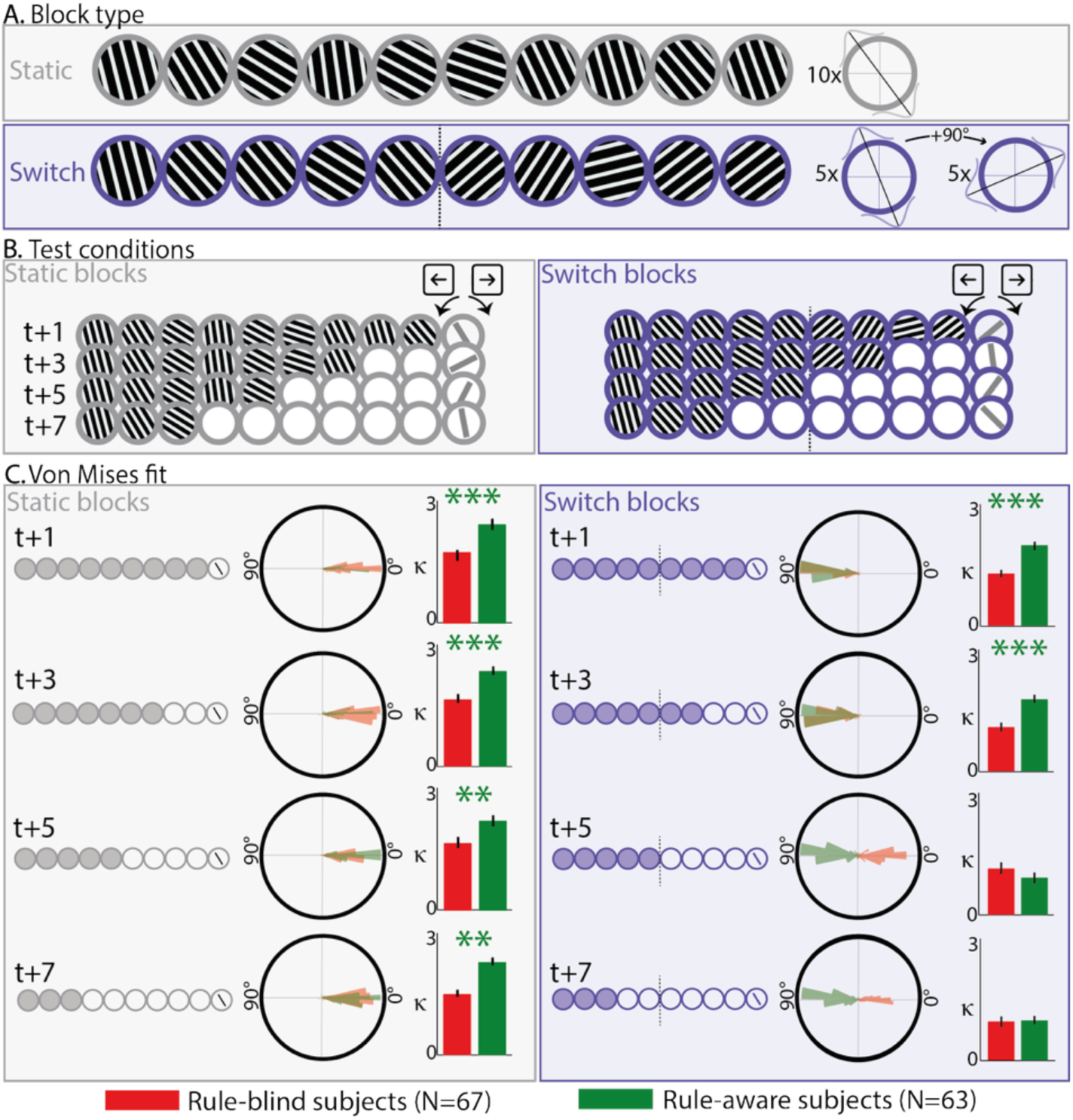
**A.** Description of the two types of blocks from Experiment 1. Each of the grating patches lasts 250ms. In static blocks, sequences were composed of 10 items drawn from a normal distribution centered on a target orientation randomly picked for each sequence. For switch blocks, the last 5 stimuli of sequences were drawn from the 90° shifted distribution compared to the first 5 stimuli. **B**. For each block, after the presentation of 6 complete sequences, participants performed different conditions, i.e., incomplete sequences of 3/5/7 or 9 first stimuli. For each, they had to predict what would have been the orientation of the last (10^th^) item of this particular sequence by rotating a bar. **C.** We compared participants’ predictions to the mean orientation before switch and fitted Von Mises distributions for each participant, condition and block. Polar plots display the histogram of the location (*θ*) of the Von Mises distribution for each participant, while bar plots display participants’ average precision (*κ*). Error bar represents the standard error of *κ* values across subjects **: ps<0.01 *** ps<0.001 (Bonferroni corrected across conditions).

Our findings, obtained from nearly 600 participants across 5 experiments, uncovered a clear bidirectional interaction between sensory integration and rule discovery. We found that participants’ timescale of sensory integration during static blocks predicted whether they would discover the latent hidden rule in switch blocks. Conversely, we found that discovering the latent rule led rule-aware participants to accurately mentally rotate presented stimuli in switch blocks before integrating them. Additional experiments controlled for general attention and working memory to confirm that the timescale of sensory integration has a unique effect on latent rule discovery. Moreover, we showed that temporal unpredictability selectively impairs rule discovery, without altering sensory integration. Finally, we used targeted manipulations of recurrent neural networks^33–35^ to establish a causal link between sensory integration timescale and latent rule discovery.

## Results

### Experiment 1: Sensory integration predicts rule discovery

We first investigated if two key human inference processes - sensory integration and rule discovery - are interacting or independent by testing if rule discovery in switch sequences could be predicted by subject performances during static sequences. First, we identified behavioral signatures diagnostic of latent rule discovery, to classify participants accordingly. Separately for each type of block (static/switch) and each type of prediction (*t*+1/*t*+3/*t*+5/*t*+7), we fitted the distribution of tilt between each participant’s response (their predicted orientation for the 10^th^ stimulus) and the mean orientation before the switch (see **Methods**). The fitted location (*θ*) provided a clear signature of rule discovery (**Fig. 1C**). Indeed, for conditions *t+5* and *t+7* in switch blocks, the orientation of the 10^th^ stimulus is shifted by an average of 90° relative to the mean of the pre-switch distribution. A participant who discovered the hidden rule should have a location *θ* close to 90°, whereas a participant who did not should have a location *θ* close to zero, as they have only seen the first half of the sequence in these conditions. Fitted location for these conditions in switch blocks revealed a clear bimodal distribution around these two behaviors (**Fig. 1C**). We split participants into two groups based on their fitted location in these conditions: *rule-aware* for participants with *θ* > 45° (N=63) and *rule-blind* for participants with *θ* < 45° (N=67, **Fig. 1C**). The order of block presentation in the experiment had no impact on the discovery of the rule (71 participants started with a static block: 48% rule-aware; 59 started with a switch block: 49% rule-aware.”).

We compared these two groups in terms of other aspects of behavior. The first step was to compare their response precision *κ* (see **Methods**). Rule-aware participants made significantly more precise predictions than rule-blind participants in static blocks (concentration *κ*: p_s_<0.01, Bonferroni corrected; **Fig. 1C**). They also made more precise *t+1* and *t+3* predictions in *switch* blocks (p_s_<0.001). This improvement in prediction precision in rule-aware participants was accompanied by a higher self-reported confidence than rule-blind participants (**Fig. S2**).

To characterize the nature of the increased precision of rule-aware participants, we used a computational modeling approach^28,29^. We decomposed each participant’s imprecisions during static blocks into two sources of internal errors: integration errors, arising from suboptimal perception and integration of successive stimuli, and response errors, reflecting suboptimal reporting of the participant’s prediction (M1; **Fig. S1**). Integration errors, but not response errors, were significantly smaller for rule-aware than rule-blind participants (rank-sum test, *p<0.001*, response error *p=0.35* **Fig. S1B**).

### Experiment 1: Sensory integration timescale predicts rule discovery

To better understand the specific sensory integration parameters which affect rule discovery in switch blocks, we derived a fine-grained model of participants’ behavior in static blocks (M2). It comprised five sources of suboptimality (**Fig. 2A**): 1/ sensory bias (γ_*sen*_) distorting stimulus orientation toward diagonals^28,36^, 2/ random noise of standard deviation σ_*int*_ corrupting the integration process, 3/ integration leak (α), capturing the timescale of sensory integration with a stronger leak corresponding to a shorter integration timescale 4/ random noise of standard deviation σ_*rep*_ corrupting the reporting of participants’ belief, and 5/ response bias (γ_*rep*_) distorting the reported orientation toward diagonals. Parameter recovery confirmed that all parameters were dissociable and interpretable (**Fig. S3**).

**Figure 2:**
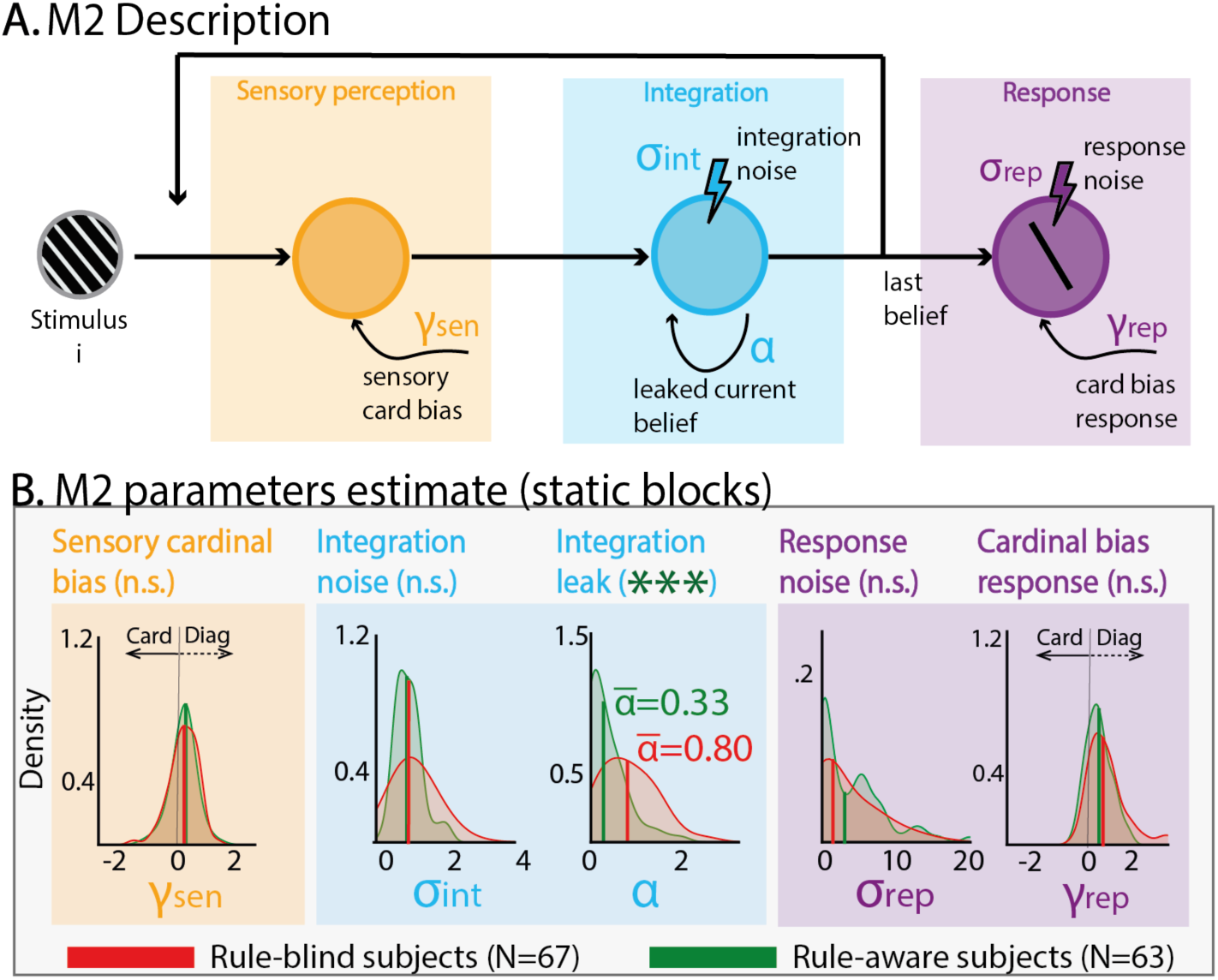
**A.** Description of Model 2 with five suboptimalities parameters (see methods). **B.** Parameter fit on static blocks and group comparison between rule-aware and rule-blind participants. ***: ps<0.001 (rank-sum test, Bonferroni corrected over the 5 parameters)

Fitting this model to participants’ behavior in static blocks revealed a selective difference in sensory integration leak between rule-aware and rule-blind participants: rule-aware participants showed lower integration leak (ie. longer integration timescale -α=0.32+/-0.05) than rule-blind participants (0.80+/-0.08; rank-sum test *p<0.001* Bonferroni corrected). No other parameter significantly differed between the two groups of participants.

### Experiments 2-3: Sensory integration timescale and rule discovery are unrelated to working memory

To test whether individual differences in working memory (WM) capacity (or more generally, in executive attention) were related to individual differences in sensory integration timescale, we carried out a first control experiment. In **Experiment 2,** a new group of participants completed static blocks (without a hidden rule) alongside an n-back working memory task using the same stimuli – by asking participants to report the orientation of a single stimulus in the sequence cued at sequence offset (see Methods). Participants’ performance in the sensory integration task and in the working memory task diverged as sequence length increased: accuracy in the n-back working memory task declined, whereas accuracy in the sensory integration task improved (ANOVA seqlen x blocktype, *p<0.001*). This performance dissociation suggests that performance in the sensory integration task does not rely strongly on working memory capacity. In line with this dissociation, individual differences in performance in the working memory task did not correlate with individual differences in the sensory integration timescale, as estimated by the integration leak α from model M2 fitted to behavior in the sensory integration task (**Fig. S4**).

To further test whether individual differences in general attention or working memory capacity were related to individual differences in rule discovery, we carried out a second control experiment. In **Experiment 3,** an additional group of participants performed *switch* blocks (with the hidden rule) together with two working memory assessments: a first assessment targeting WM capacity, and a second assessment targeting WM maintenance. Although participants’ performance in the two WM assessments correlated with each other (Spearman correlation R=0.31 *p<0.01*), neither distinguished rule-aware from rule-blind participants in the switch blocks (rank-sum tests both *ps>0.24*, **Fig. S4**). Together, these results indicate that individual differences in working memory performance do not explain the observed effect of sensory integration timescale (reflected in the integration leak) on rule discovery.

### Experiment 1: Rule discovery alters sensory integration

Analysis of the static blocks provided clear evidence for a “bottom-up” interaction between sensory integration timescale (as measured by participants’ leak) and rule discovery. To explore a possible “top-down” interaction of rule discovery on human integration strategy, we turned to the analysis of participants’ behavior in switch blocks. We compared two hypotheses: Either participants that discovered the rule do not change their integration process but only add a correction offset to their response to account for the rule (Model 3), or they flexibly apply mental rotation to each of the first 5 stimuli, prior to sensory integration (Model 4).

We fitted those two models (**Fig. 3A**) on switch data together with the baseline model without any offset (M2; **Fig. 2A**). Model comparison revealed the integration offset model (M4) to be the best fit for rule-aware participants (AIC: M2=671, M3=647, M4=523; *p<0.001*, **Fig. 3A**). We checked that the offset rule implemented in this model (same offset for the first five stimuli and no offset for the last five) was correct by fitting a complementary model with 9 offsets, one for each stimulus (Model 5). This confirmed our hypothesis, showing rule-aware participants had an exact representation of the switch rule and applied a 90° offset to the first 5 stimuli and no offset to the last 5 (**Fig. S5**). Discovering the rule modified participant’s sensory integration process, as they integrate mentally rotated information, rather than simply accounting for the rule in their reporting process.

**Figure 3:**
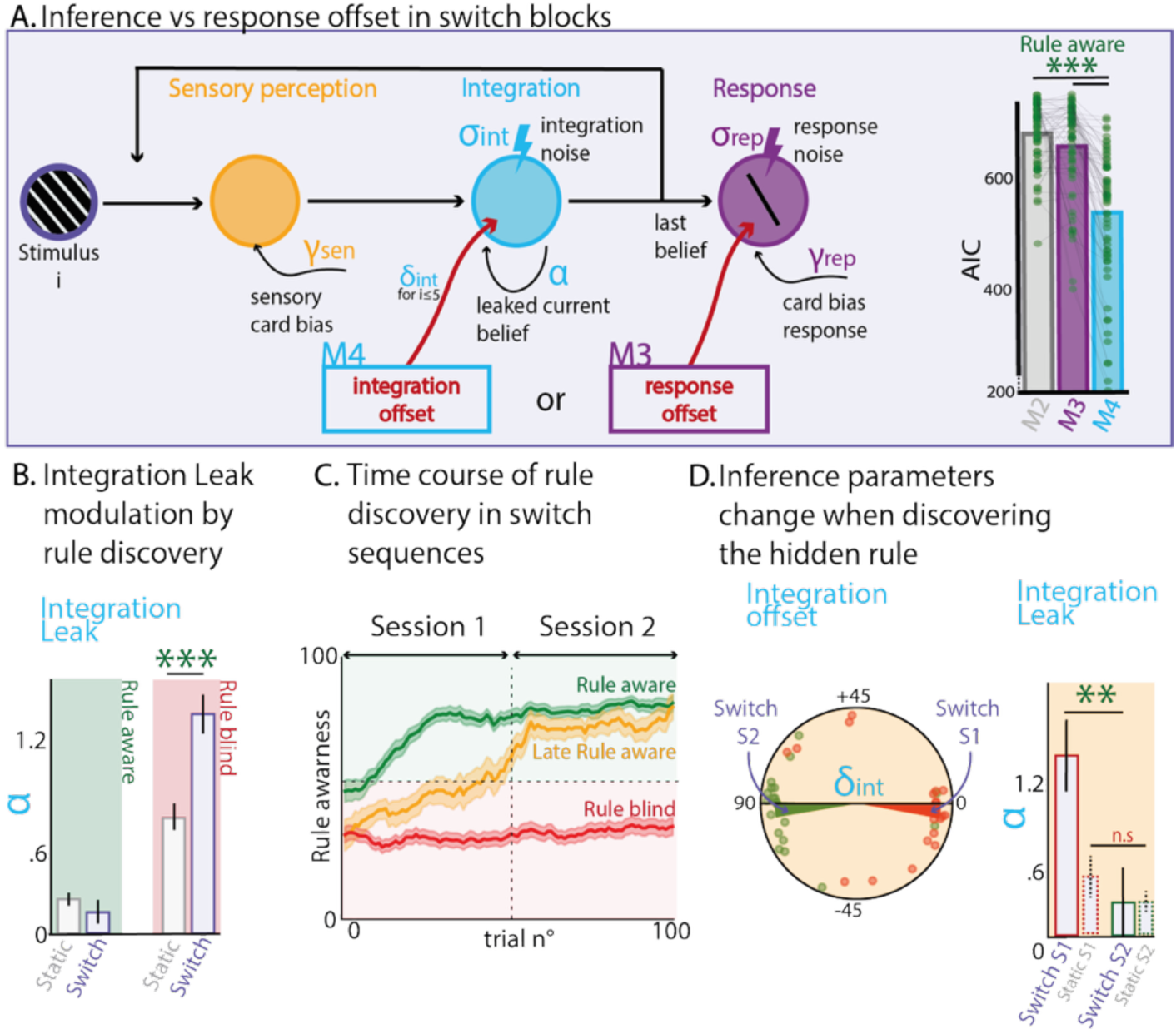
**A. Left.** Description of Models 3 and 4. Model 3 is similar to Model 2 (fig. 2) with an additional offset parameter applied to the response while Model 4 has instead an offset parameter applied during the integration of the five first stimuli of the sequence. **Right.** AIC comparison when fitted on rule-aware participants’ responses to switch blocks for Models 2 (‘no offset’), 3 (‘response offset’), and 4 (‘integration offset’). **B.** Comparison of the integration leak within participants, between static and switch blocks, for rule-aware and rule-blind participants. ***: ps<0.001 (rank-sum test). **C.** Time-course of the rule discovery for each group of subjects. The y-axis represents the probability that a given trial was closer to the rule-aware answer compared to the rule-blind answer. **D.** Analysis of the 20 late rule-aware participants that were rule-blind in session 1 but rule aware in session 2. Discovering the rule in session 2 modified their integration by adding a 90° offset but most importantly drastically reducing the leak, further evidencing the causal role of rule discovery on integration timescale.

To further support this idea, we explored one of its predictable implications. Participants who did not discover the rule should perceive switch blocks as more volatile than static blocks, while participants who found the rule can use this knowledge to explain away perceived volatility. As perceived volatility is typically accompanied by a smaller integration timescale^37,38^, we compared the leak during the static and the switch blocks for both groups. We found an increased leak in switch compared to static blocks for rule-blind participants (α switch=1.5+/-0.13, static=0.79+/-0.08; rank-sum test *p<0.001*), up to the point where they almost ignored pre-switch stimuli, while rule-aware participants kept a similar leak, revealing the use of the rule knowledge to reduce perceived volatility of switch sequences (α switch=0.31+/-0.06, static=0.32+/-0.05; *p=0.6*; **Fig. 3B**, interaction *p<0.001*). Integration noise was also modulated by rule discovery, in a way that suggests that the mental rotation applied to stimuli in switch blocks by rule-aware participants is associated with an additional source of random errors (**Fig S6**).

To investigate whether discovering the rule causally influences changes in integration parameters (rotation and leak), we performed a quasi-causal analysis by focusing on a specific subgroup of participants, labeled as *late rule-aware (N = 20)*: classified as rule-blind in the first session (on day 1) but rule-aware in the second session (on day 2). If rule discovery genuinely modifies sensory integration parameters, we expected to observe changes in parameter values in the switch block of the second session compared to the first session. Trivially, rule-aware participants had a greater integration offset but more importantly a substantial reduction of integration leak in the second session compared to the first session, only during *switch* blocks (*p<0 .01* - **Fig. 3C**).

### Experiments 4-5: Temporal prediction selectively affects rule discovery

Sensory integration and rule discovery are not the only two forms of inference humans are known to be capable to perform. Indeed, presented with stimulus sequences, humans can, not only predict *what* they expect to occur but also *when* they expect a given element to appear. To investigate if temporal prediction affects the two others, we designed a jittered version of Experiment 1 and measured the influence of temporal unpredictability on both sensory integration and rule discovery. We reasoned that if sensory integration and rule discovery reflect distinct forms of inference – rather than two aspects of a single core cognitive ability^24^, temporal unpredictability could selectively disrupt only one of these forms of inference (Experiment 4, see Methods).

We first replicated all main effects of Experiment 1 (**Fig S6**). Then, we tested the influence of temporal jitter on sensory integration by fitting Model 2 to the data from static blocks with temporally jittered sequences (Experiment 4). We found no difference in the fitted parameters compared to those from static blocks with rhythmic sequences (Experiment 1) (rank-sum test all *ps>0.05*; **Fig 4A**). We confirmed this absence of difference in sensory integration parameters between rhythmic and temporally jittered sequences using a within-subject design (Experiment 5), where the same participants performed static blocks with and without temporal stochasticity (**Fig. S7**).

**Figure 4:**
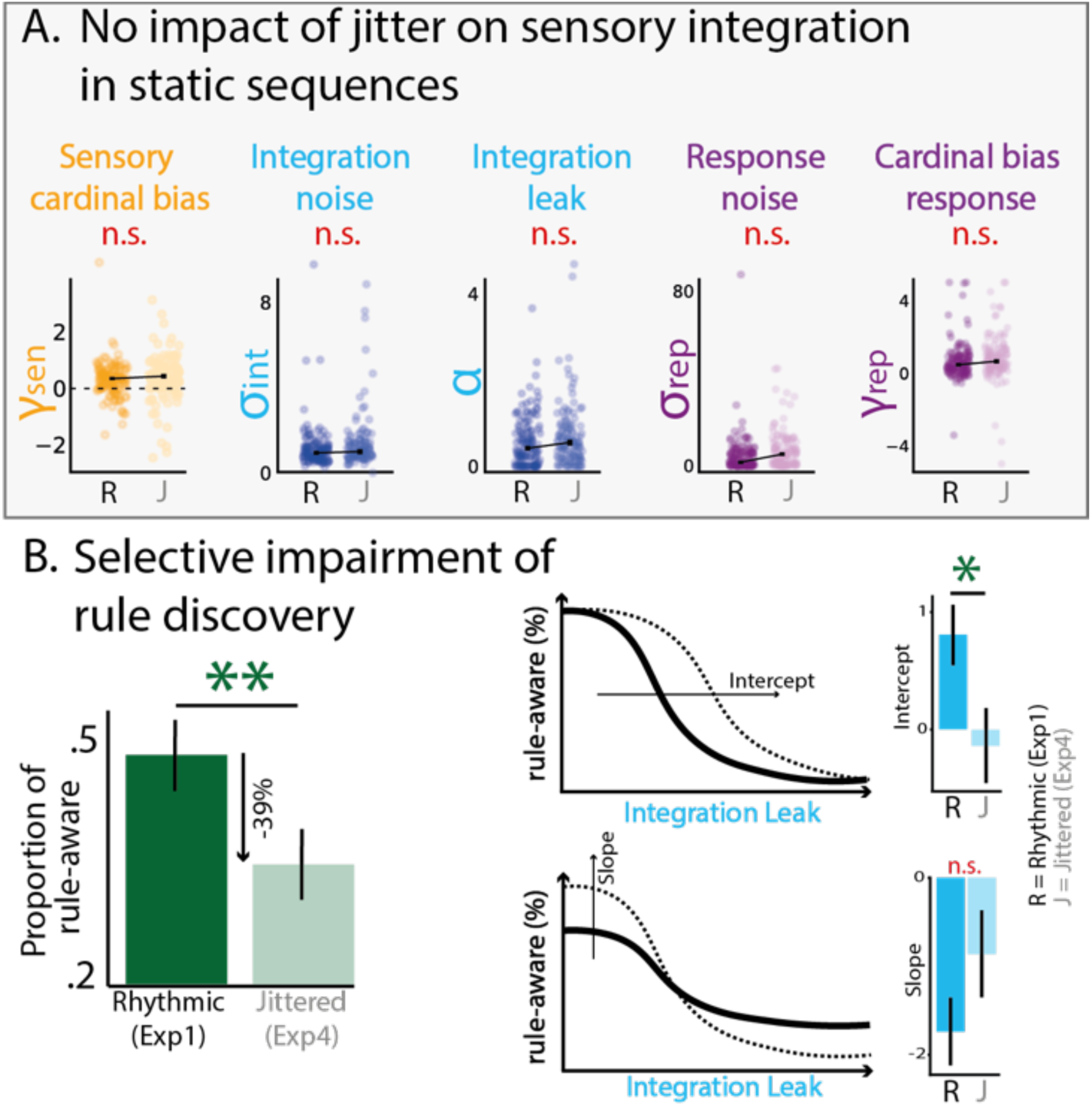
**A.** Comparing integration parameters during static sequence for rhythmic (Experiment 1) and jittered (Experiment4) presentation revealed no impact of temporal stochasticity on integration. **B.** However, jitter during the presentation significantly reduced the percentage of participants that found the hidden rule in switch sequences (p=0.01). Fitting sigmoid between leak parameter during static sequences and rule-awareness probability gave a different intercept (p<0.05) but a similar slope, revealing independent impact of jitter and leak on rule discovery.

Despite these unchanged integration parameters, temporal stochasticity significantly reduced participants’ likelihood of discovering the rule during switch blocks (−39%, *p=0.01*, bootstrap resampling; **Fig. 4B**), revealing a specific influence of temporal prediction on rule discovery. We found that this decreased probability of discovering the latent rule with temporally jittered sequences (in switch blocks) was independent of individual differences in integration leak (in static blocks) by fitting a logistic function that relates integration leak to the probability of rule discovery in each experiment. The intercept term, but not the slope, was significantly higher for rhythmic sequences (Experiment 1) compared to temporally jittered sequences (Experiment 4, bootstrap resampling *p<0.05*), indicating that the effect of temporal stochasticity on rule discovery is not mediated by changes in integration leak (**Fig. 4B**; see **Fig. S8** for additional control analyses).

### RNN simulations: Shorter sensory integration timescales prevent rule discovery

Previous analyses showed that participants’ integration timescale (measured by integration leak) was specifically associated with their capacity to discover the hidden rule in switch blocks. Detailed analyses of late rule-aware subjects (Exp 1) and jittered experiments (Exp 4&5) gave insights regarding the causal nature of the effect of sensory integration timescale on rule discovery. To further strengthen these findings, we turned to recurrent neural networks (RNNs) to test directly for a causal effect of sensory integration timescale on latent rule discovery.

In practice, we trained RNNs to predict each upcoming stimulus in the sequence. This approach allowed us to measure the network’s surprise at each stimulus in the sequence (see **Methods**). Based on these surprise estimates, we defined a *Rule Learning Index (RLI)* as the network’s ‘surprise’ to the switch (stimulus 6), relative to the average surprise across all other stimuli. This index, unavailable from human behavior, captures the latent dynamics of the ability of the network to correctly predict the switch rule and thus provides a quantitative metric of rule discovery in RNNs. In particular, it allows for precise dissection of the learning of sensory integration and rule discovery. Throughout the training, we observed a rapid decline in surprise for all stimuli except the switch, corresponding to the learning of pure sensory integration. This was followed by a later decrease of the surprise evoked by the 6^th^ stimulus, indicating rule discovery (**Fig. 5B**). This dynamic is revealed in a sharp initial rise in the RLI (sensory integration learning), followed by a slower decrease (rule learning) until convergence.

**Figure 5:**
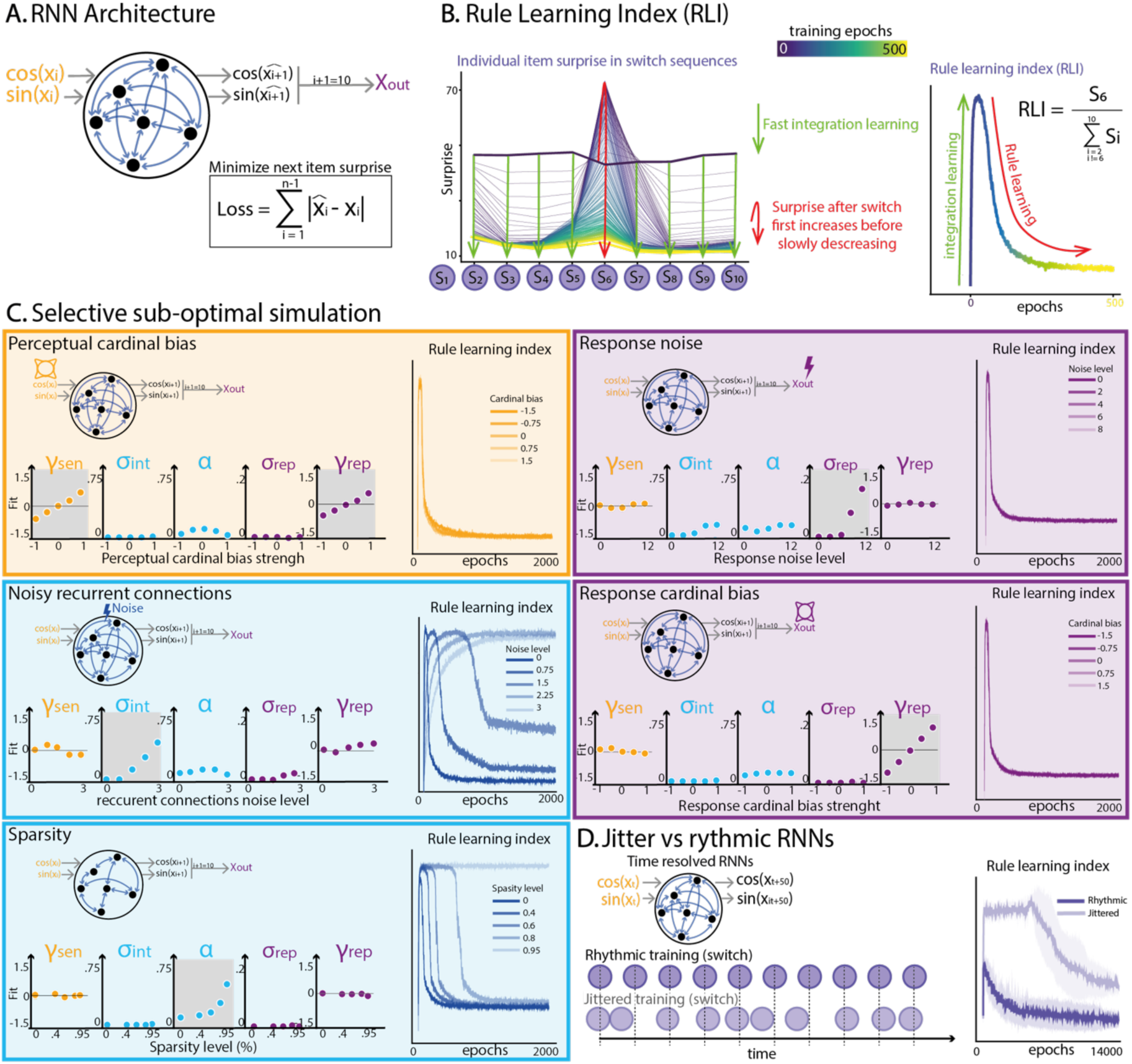
**A.** Architecture of the Recurrent Neural Network (RNN). The RNN receives as input the current stimulus orientation (x_i_), and is trained to predict the next orientation (x_i+1_). The difference between the predicted and actual orientation defines the ‘surprise’. **B. Left:** Surprise for each stimulus (2-10) in the switch blocks as a function of training epochs (color-coded from purple to yellow). Green arrows highlight the rapid decrease in surprise for stimuli 2/3/4/5 and 7/8/9/10 corresponding to the learning of sensory integration predictions. The red arrow highlights the slower decrease in surprise following the switch, corresponding to the discovery of the rule. **Right:** This learning dynamic is summarized by the Rule Learning Index (RLI), defined as the ratio of surprises related to rule learning (stimulus 6) and sensory integration (stimuli 2-to-5 and 7-to-10). **C**. Five variants of the RNN designed to implement each of the five suboptimalities defined in Model 2, with five levels of suboptimalities (from low to high) for each. 1. Perceptual cardinal bias applied to each orientation xi. 2. Integration noise modeled via noisy computation in the recurrent layer. 3. Integration leak obtained by sparsifying to the recurrent layer. 4&5. Response-related parameters were applied at the output level (Xout). **Bottom graphs on each panel.** After training, we fitted Model 2 to each RNN’s predictions on static blocks to estimate how each manipulation impacted each of the parameters. **Right graph on each panel.** For each level of suboptimality, we estimated the RLI to evaluate how it affected the learning dynamic.

To causally explore the role of each suboptimality, we aimed at selectively modulate each in our networks, and observe its effect on RLI. For that, we aimed at introducing five selective perturbations that trigger the five suboptimalities observed in human behavior (**Fig. 2**). We simulated the behavior of selectively perturbed RNNs and fitted Model 2 to the RNN behavior during static blocks – similarly to what was done in humans to estimate their suboptimality parameters. We showed that distorting the network input with cardinal bias triggered a sensory cardinal suboptimality in RNN behavior. Similarly, distorting RNN response with cardinal bias or random noise introduced respectively cardinal bias response and response noise in their behavior. Most importantly, adding noisy recurrent connections triggered integration noise (σ_*int*_) and sparsifying the hidden layer specifically triggered leak (α) in RNN behavior (**Fig 5C**).

We used the *RLI* to assess how each suboptimality causally affects the learning trajectory in switch sequences and showed that, integration noise and integration leak modified RLI dynamics. More precisely, while adding integration noise (σ_*int*_) via noisier recurrent connections globally impaired both sensory integration and rule discovery, increasing integration leak (α) specifically delayed discovery of the switch rule without altering sensory integration (**Fig. 5C**). We considered learning timecourse as a complementary metric to closely match humans’ analyses (**Fig3C**) and replicated our main effects (**Fig S11**). We also confirmed that this relationship was not limited to the particular case of a single switch in the middle of the sequence by replicating this analysis using another rule with sequences of the form AABBAABBAA (**Fig. S9**).

To support the selective interaction between temporal predictability and rule discovery observed in Experiments 4-5, we trained a new set of RNNs to make temporally resolved predictions and trained them with rhythmic or temporally jittered stimulus sequences, using the exact same parameters used in Experiment 4. A Rule Learning Index (RLI) adapted to this temporally resolved condition showed that, like humans, the training of RNNs to perform sensory integration was unaffected by temporal stochasticity, whereas the latency at which the same RNNs discovered of the latent rule was strongly delayed by temporal stochasticity (**Fig 5D**, see details in **Fig S10**).

## Discussion

To make sense of sequences of stimuli, humans rely on multiple inferences often studied independently, leaving unknown how they interact to form a comprehensive learning system in humans. A few recent studies have begun to address this issue, by investigating the impact of temporal prediction on statistical learning^39^, or by asking human participants to classify sequences of stimuli drawn either from deterministic rules or from probabilistic distributions^40^. The present study aims at understanding how sensory integration, temporal prediction, and rule discovery interact to form a flexible inference system that is able to make sense of sensory events unfolding in time. Our results revealed a strong two-way interplay between sensory integration and rule discovery: (1) a bottleneck exerted by the integration timescale (leak) of sensory integration on the probability of further discovering the hidden rule, and (2) the application of the hidden rule to incoming stimuli prior to their integration, as well as the use of the rule knowledge to adapt integration timescale. Our findings further reveal a selective impact of temporal prediction on rule discovery, leaving sensory integration unaffected.

First, Experiment 1 showed that participants’ integration timescale (as measured by integration leak^28,30,41,42^), measured in rule-free sequences, predicts the likelihood of discovering a rule in other sequences (**Fig. 2B**). Note that the hidden rule in switch sequences triggered not a gradual but a brutal 90° shift in mean stimulus orientation between the 5^th^ and 6^th^ stimuli, which therefore did not theoretically require a long integration timescale to be detected. Participants who did not find the hidden rule in switch blocks showed moderate, not extreme integration leak in static blocks. As integration timescale likely reflects one’s prior belief about environmental volatility, participants who expect a highly volatile environment lower their integration timescale – making them less likely to detect the latent rule. On the other hand, participants who expect a low environmental volatility rely on a larger integration timescale and can look for a latent rule explaining away the change in statistics from the middle of sequences. Importantly, we verified that this effect of sensory integration timescale on rule discovery is not attributable to differences in general attention or working memory and was robust to which block type was first experienced by subjects.

To assess the possible causal role of integration timescale in participants rule-blindness, we applied selective causal perturbations to RNNs trained to perform the same task as human participants. This effort is part of a growing body of research that leverages these flexible artificial systems to understand cognition, not by fitting their free parameters (connection weights) to behavior^43,44^, but by analyzing the structural and functional configurations which, through constrained optimization, result in the same suboptimalities as biological brains^45–47^. Sparsifying recurrent connections produced a graded increase in integration leak during static blocks without affecting any other suboptimality. In contrast, adding computational noise in recurrent units^29,45^ increased integration noise independently of leak. In switch blocks, these manipulations revealed a clear distinction: increasing sparsity impaired rule discovery without disrupting the training dynamics of sensory integration. In contrast, adding computation noise in the hidden layer (resulting in increased integration noise) impaired both the training of sensory integration and the latency of rule discovery. This distinction was captured in RNN surprise dynamics during training: networks first reduced surprise to all stimuli except the 6th (indicating sensory integration), then gradually to the 6th (indicating rule discovery). Only integration leak delayed the latter without altering the former. The link between leak and delayed rule discovery generalized to sequences with a different hidden rule structure (with 90° shifts every two stimuli), supporting the generality of this causal relationship. Although the number of trials required to train humans and ‘blank-slate’ RNNs on our task differs by orders of magnitude, the shared impact of sensory integration timescale on rule discovery similarly constrains humans and RNNs.

While integration timescale was reliably and consistently tied to poorer rule discovery abilities both in humans and RNNs, our results also show that discovering the hidden rule modified how human participants integrate stimuli in switch blocks. In humans, discovering the rule causally triggered the mental rotation of individual stimuli prior to integrating them, in order to predict the final (10^th^) stimulus. This finding directly contradicts dominant theories and computational models of sensory integration, which typically postulate that integration occurs within the ‘native sensory space’ defined by stimulus features (e.g., orientation)^48–52^. If sensory integration was encapsulated in this way, rule discovery would result in a post-hoc shift applied to the integrated result. This is not what we observed: participants’ behavior was best explained by a model in which stimuli were mentally rotated according to the hidden rule during integration. Such abstract mental manipulations of stimuli have already been described in rule learning tasks, notably in the context of temporal rule in stimulus sequences, such as the reordering^21,23^, mapping^53^ or compression^54^. These findings are much sparser in the context of noisy sensory integration. Our findings extend previous results showing dissociated neural correlates of the sensory processing of individual stimuli and the integration of their associated category evidence^9,11,55^.

Moreover, the discovery of the latent rule also protected rule-aware participants from the typical decrease in integration timescale that often accompanies increased volatility^37,38^. Indeed, despite a much larger sensory volatility in switch blocks compared to static blocks, the knowledge of the rule allowed rule-aware participants to explain away this sensory volatility and therefore keep their integration timescale as long as in static blocks. By contrast, rule-blind participants perceived switch blocks as more volatile, and therefore – and adaptively so – reduced drastically their integration timescale reflected by a much larger integration leak. These findings further show that rule-blind participants were not insensitive to the difference between static and switch blocks. Rather, they interpreted the difference between blocks as an increased sensory volatility in switch blocks, instead of a latent rule.

Our study is not the first to investigate the impact of hidden rule discovery on decision making. However, the hidden rules used in previous studies typically did not interact with the processing of presented stimuli. For example, the hidden rule used in Schuck et al^56^ afforded bypassing entirely the processing of the pattern of dots to focus solely on their color to decide. This bypass prevented testing for the interplay between stimulus processing and rule discovery. By contrast, in our task, a leak in sensory integration directly prevented participants from discovering the hidden rule, an objective limitation in constructing accurate predictions of upcoming stimuli. RNN simulations showed that this effect was not limited to the specific AAAAABBBBB rule used in our task but also generalized to other rules like AABBAABBAA, even though future research in humans should investigate the set of rules over which this effect generalizes. The rule we have used – with a switch in the middle of a sequence of stimuli – corresponds to the “reversal learning” paradigm that has been extensively used to study adaptive learning in humans and animals^57–61^. We have adapted this paradigm so that the switch followed a hidden rule, which could either be perceived as predictable (if the hidden rule has been discovered) or as randomness in stimulus sequences (if the hidden rule has not been discovered).

RNN perturbations showed that integration leak selectively emerged only for extremely sparse networks, resulting in a substantial decrease in the number of computations performed. Indeed, RNNs with only 5% of active connections showed an integration leak comparable to human participants, without any other suboptimality. The integration leak measured in human participants could therefore partly stem from a cost-benefit optimization, which trades the benefits of efficient sensory integration and rule discovery against the cost of more dense computations. Such cost-benefit arbitration has been postulated to explain other suboptimalities in decision-making such as integration noise, which could arise from the flexibility of associative circuits that can map arbitrary functions of sensory features (here, the integration of stimulus orientations with rule-predicted shifts) onto actions^29,45^. Indeed, the attentional selection and pooling of sensory signals according to a current context^62,63^ by low-rank associative neural circuits has been theorized to be critical for decision accuracy^64^.

Finally, we found that temporal predictability (i.e. predicting the timing rather than the content of a perception^65^), was selectively interacting with rule discovery, while having no direct impact on sensory integration. This dissociation suggests that the precise timing at which the latent switch occurred mattered for triggering latent rule discovery, possibly because temporal jitter triggered stronger surprise for all stimuli of the sequence, therefore reducing the relative importance of the surprise after the switch (6^th^ element) compared to the surprise induced by other stimuli of the sequence. By contrast, sensory integration of high-contrast stimuli seems unaffected by temporal jitter – consistent with earlier work showing its robustness to temporal parameters of sequences^66^. Together, our findings emphasize that human inferential abilities result from multiple processes with complex interdependences (**Fig 6**). It further highlights the need to conjointly study these several inference abilities to obtain a clear picture of how humans are able to build a rich mental representation of uncertain environments.

**Figure 6:**
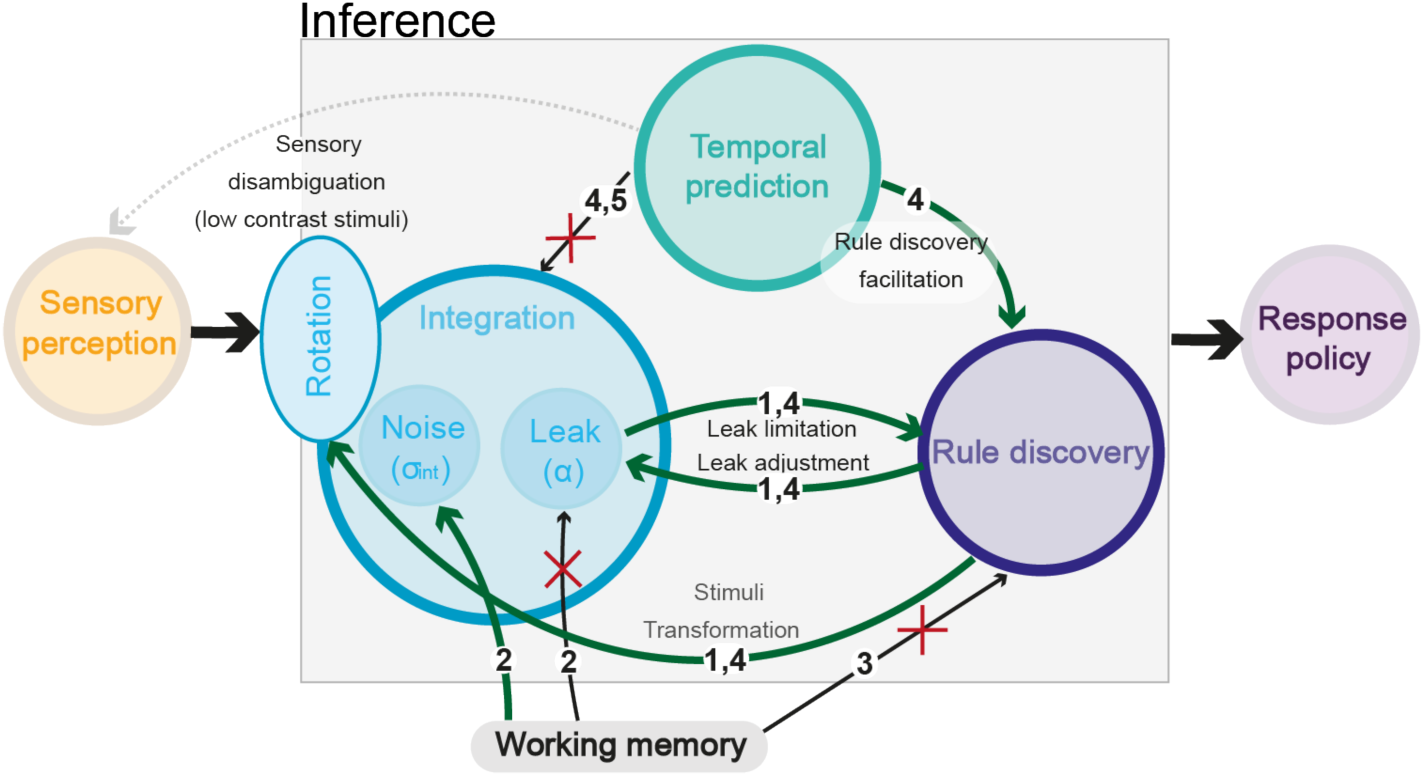
Visual summary of our findings across all experiments: green arrows represent demonstrated links between cognitive abilities and crossed black arrows where no such links were found. Numbers correspond to experiments testing each arrow. The grey arrow between temporal prediction and sensory perception corresponds to a known effect of sensory disambiguation, not shown here.

## Materials and Methods

### Participants

All participants for all experiments were UK based adults (>18yo), recruited online via the Prolific platform and gave informed consent and the study was approved by Inserm Ethical Review Committee (IRB00003888 on 13/11/2018). *Experiment 1:* 130 participants performed two sessions of the task on consecutive days (52 female, mean age = 41,5). They were paid 8£ for the first and 9£ for the second session. *Experiment 2:* 100 participants performed the task (53 female, mean age = 42.3) and performed a single session retributed 7£*. Experiment 3:* 102 participants were recruited online (49 female, mean age = 41.6) and performed a single session retributed 7£*. Experiment 4:* 123 participants performed two sessions of the task on consecutive days (72 female, mean age = 41.1). They were paid 8£ for the first and 9£ for the second session. *Experiment 5:* 126 participants performed two sessions of the task on consecutive days (57 female, mean age = 38.7). They were paid 8£ for the first and 9£ for the second session.

### Experimental paradigm and stimuli

All experiments were coded in JavaScript using jspsych toolbox^67^.

**Experiments 1&4** consists two types of blocks: in static blocks, each stimulus of the sequence is a grating patch whose orientation is drawn from a normal distribution (σ = 15°). In *switch* blocks, we added a 90° rotation from the middle of the sequences. Importantly the means of the distributions were randomly chosen for each trial so that the abstract +90° rule must be learnt. Each sequence was sequentially presented at 2Hz, rhythmically in Experiment 1, or with random ISI (uniform [0, 500] ms) for Experiments 4 and 5. The block type was indicated by the color of the circle surrounding the stimuli. Block order was random across participant. In each block, after the presentation of 6 full training sequences, participants were presented 20 with incomplete sequences, stopping after 3, 5, 7 or 9 stimuli (5 of each), and had to predict the orientation of the last (10^th^) stimuli by rotating a bar, and estimate their confidence level. 4 full sequences were randomly placed between the incomplete test sequence to consolidate participant learning. After each answer, they saw colored feedback indicating how far they are from the real orientation. They performed a total of 400 test trials over the two sessions.

**Experiment 5** had an identical procedure to Experiments 1 and 4 but with only static trials, half of the blocks being rhythmic, and the other half jittered.

**Experiment 2** had participants perform static blocks, identical to Experiment 1 followed by memory blocks in which they were presented with sequences of stimuli and asked on each trial to report the orientation of one of the stimuli in the sequence (selected randomly on each trial). In both conditions, sequences could contain 3, 5, 7 or 9 oriented stimuli. Orientations were drawn uniformly in working memory blocks so that the orientation of a stimulus could not be predicted (by integration) from the orientation of another stimulus better than chance.

**Experiment 3** had participants perform sensory integration blocks with the same latent rule as in Experiment 1 (Switch blocks) and two types of working memory blocks measuring their ability to memorize the orientation of stimuli at positions 1-5 in the sequence (working memory capacity block) or their ability to maintain the orientation of a cued stimulus at positions 1-5 in the sequence (working memory maintenance block).

### Preregistration of Experiment 1

The procedure, as well as the primary outcome of this analysis (bidirectional interactions between sensory integration and rule discovery), has been pre-registered prior to data collection. The relevant details can be found here: https://aspredicted.org/wy43v.pdf

### Von Mises distributions fit

We performed model free analysis by fitting von Mises distributions of the difference between participants’ answers and the average orientation of the five first stimuli (only three first for t+7 conditions) of the sequence for each condition separately. Von Mises distribution have two parameters: location (*θ*) which indicates the average direction of the distribution, and the concentration (*κ*), which reflects variability of the response.

### Model design and fit

We performed model-based analyses with several models:

**Model 1**: see details in **Fig. S1**

**Model 2**: This second model comprises 5 parameters. Sensory cardinal bias (γ_*sen*_), integration noise (σ_*int*_), integration leak (α), response noise (σ_*rep*_) and response cardinal bias (γ_*rep*_). σ_*int*_ and σ_*rep*_ were modeled like in Model 1. For the sensory cardinal bias, each orientation of the sequence was modified through the function 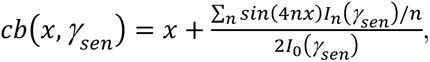 with *x* the orientation, γ_*sen*_ the bias strength and *In* the Bessel function of order n. For the response cardinal bias, the same function was applied on the response *cb* (*resp*, γ_*rep*_). The leak was modeled with an exponential decrease affecting the weight attributed to each stimulus in the participant response^38,55,68^ *–* 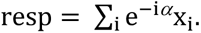 Integration noise was drawn from a normal distribution of variance σ_*int*_ and added to each element before summing. Response noise was drawn from a normal distribution of variance σ_*rep*_ and added to the response. Models 1 and 2 were fitted to static blocks to estimate participants’ parameters when performing a sensory integration task.

**Model 3**: Model 3 is equivalent to Model 2, but we added a parameter modeling a possible fixed offset that participants could apply to all answers. resp = resp + offset

**Model 4**: Model 4 is equivalent to Model 2 but with a fixed offset applied to the five first orientation of each sequence. – 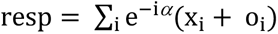 with o_i_ = offset for i ≤ 5 and 0 for i > 5.

**Model 5**: Variation of Model 4 presented in SI, were each of the 9 stimuli of the sequence can have a different offset. – 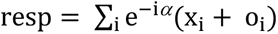 with o_i_different for each i value

Like in previous studies^69,70^, we fitted each model using BADS^71^ toolbox in MATLAB with 10,000 bootstrap resamples to minimize the negative log-likelihood and find the best fitting parameters. Models 1 and 2 were fitted on data from static blocks only while Models 3,4 & 5 were fitted on data from switch blocks. We assessed for statistical significance between parameters models using rank-sum test given the non-normality some parameter distributions and applied Bonferroni correction. We validated our model and the fitting procedure using recovery and simulations (**Fig S3**)

### RNN simulations

We first designed a simple RNN composed of a 64×64 recurrent layer. The input and output orientations are coded by a two-dimensional array with cos and sin. Importantly, the loss of the RNN was chosen to mimic human like computation. Indeed, when given an input, the networks predict the orientation of the next stimulus. The absolute value of the difference between the prediction and the actual orientation is called the ‘surprise’ and the loss of the sequence is the sum of the surprises in the sequence. That way, networks are trained with a human-like strategy.

We then implemented each suboptimalities in the RNN. The sensory cardinal bias was introduced by applying a cardinal bias for each stimulus of the sequence before the input (see Model design for the implementation of this bias). We used five cardinal bias values: [-1, -0.5, 0, 0.5, 1]. To implement the leak, we made the recurrent layer sparse by fixing a given proportion of the connections to zero. This proportion was fixed to 0, 40%, 60%, 80% or 95%. To implement integration noise, we added random noise in the computation of the recurrent layer activity from a centered normal distribution with a std of 0, 0.75, 1.5, 2.25 or 3. The two response parameters did not modify the RNN per se, but were applied on Xout: either random noise or cardinal bias. To test if those modifications were specifically affecting only one parameter of the sensory learning mechanism, we trained 20 randomly initialized RNNs on 2000 epochs of 32 complete static sequences, for each parameter value.

We then simulated the behavior after training on an occurrence of our experiment (for comparison purposes between the RNNs, we always used the same 20 task occurrences that correspond to the task presented to the first 20 participants of Experiment 1). We could therefore fit the simulation results of the static sequences with Model 2 and plot the impact of the manipulation on the fitted parameters, averaged across the 20 RNNs (**Fig. 5C**). We explored the impact of these modifications by plotting a summary metric we called *Rule Learning Index* (RLI) capturing the dynamic of the integration and rule learning through training. This metric was computed as the ratio of the surprise elicited by the 6^th^ stimulus and the average surprise elicited by all other stimuli (**Fig. 4B**). For methods on RNNs adapted to time resolved inputs, see **Fig S10**.

## Acknowledgments

This work is supported by a postdoctoral grant Espoir de la recherche from the Fondation pour la Recherche Médicale (FRM) awarded to L.B (SPF202309017516), a grant from the French National Research Agency (MONODEC, ANR-23-CE37-0028) awarded to V.W. as well as a European grant (ERC, SPEEDY, ERC-CoG-101043344) awarded to B.M.

## Supplementary Information

**Figure S1:**
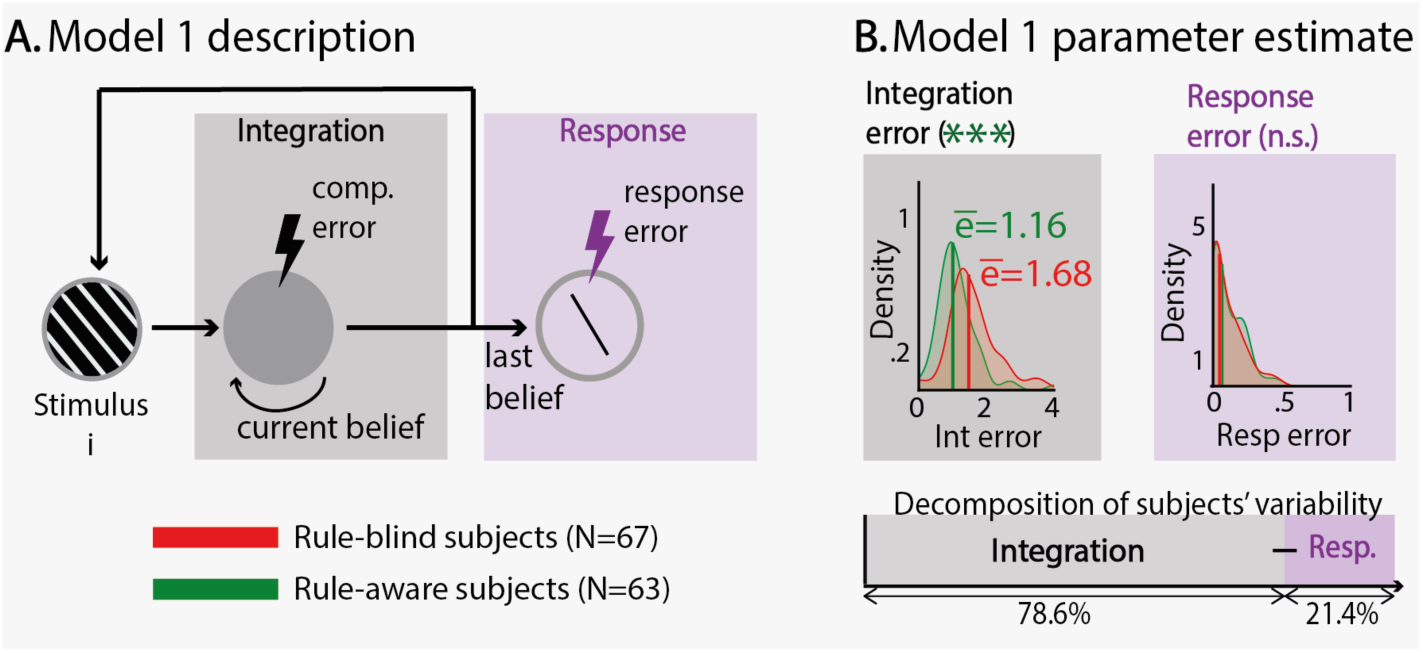
Model 1 dissociating computational from response errors. This model performs integration of evidence with two sources of uncertainties: computational and response errors. The first is modelled by a random error for each orientation drawn from a centered normal distribution with a variance σ_*inf*_. The latter is modelled by a random error on the final response with variance σ_*rep*_. To estimate the impact of each error source on the overall variance, we fitted this model to estimate σ_*inf*_ & σ_*rep*_ for each participant and simulated the response with σ_*inf*_ only, σ_*rep*_ only or both. We computed the average error for those simulations. The variance attributed to σ_*inf*_ is the ratio between the average error of the simulation with σ_*inf*_ and the average error with both sources.

**Figure S2:**
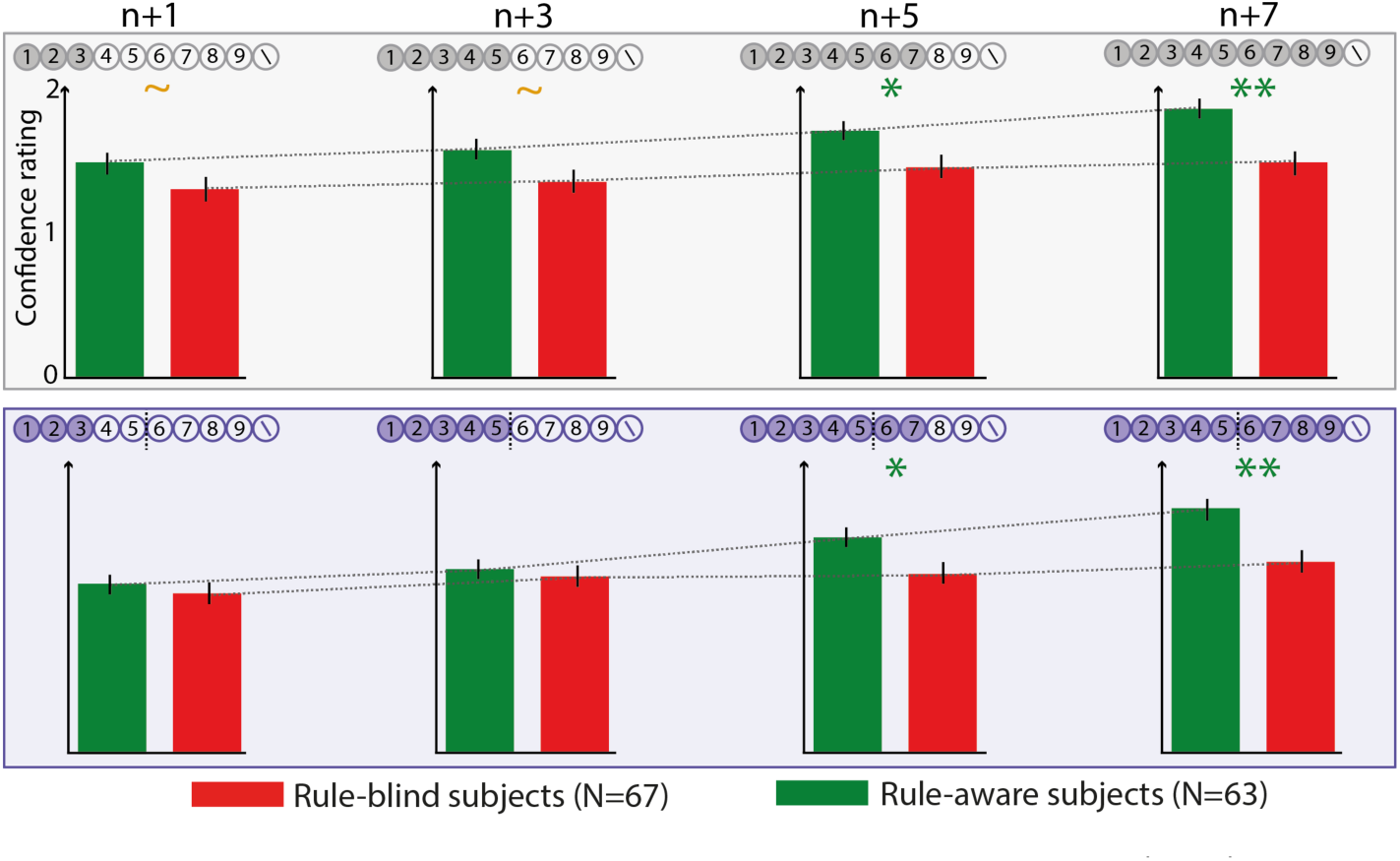
Confidence judgments for each condition, split between participants who found the rule (green) and participants who did not (red). Congruently with their small integration leak, participants who found the rule increase their confidence with sequence length (they integrate more stimuli).

**Figure S3:**
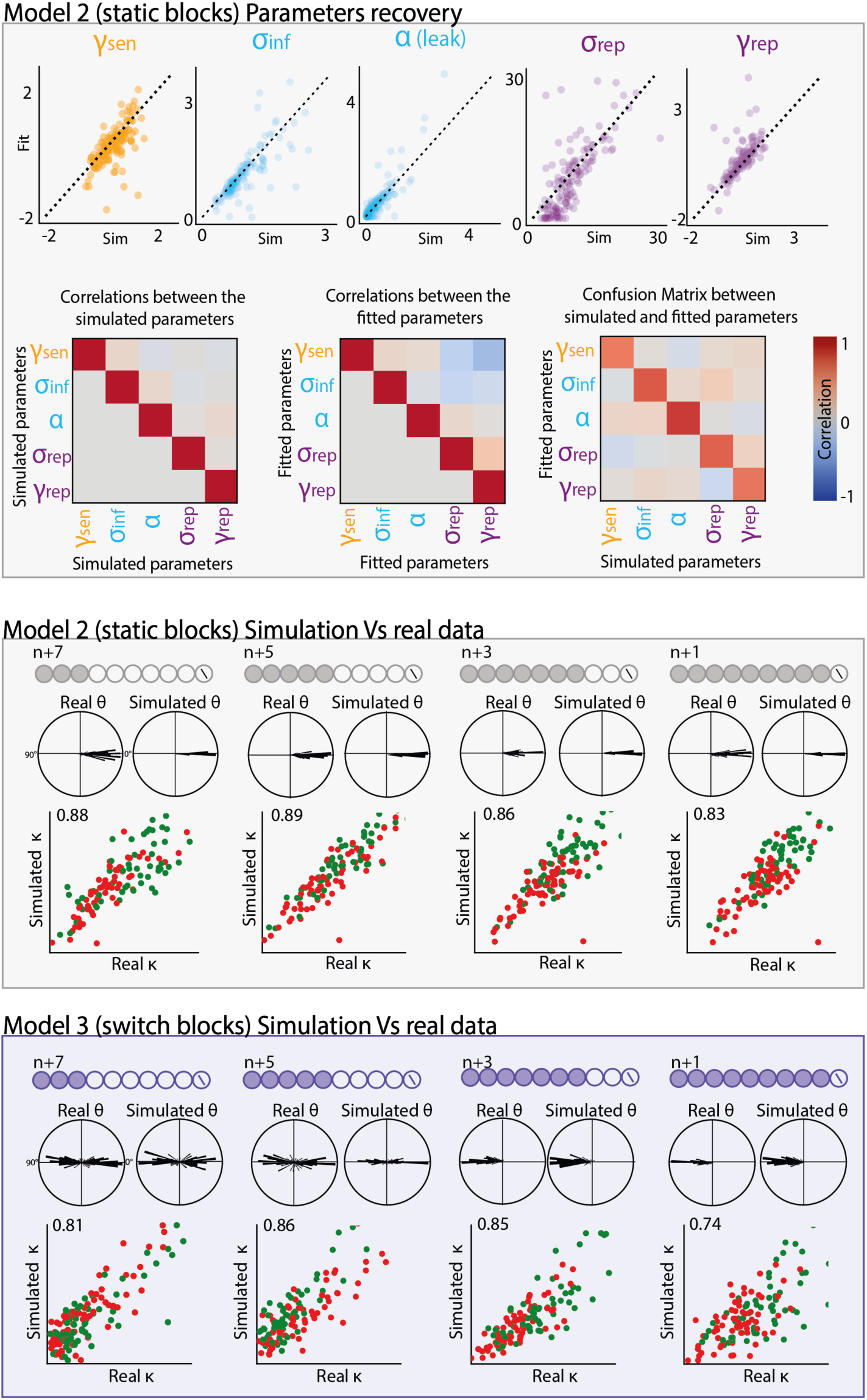
Validation of the different models. **A.** To ensure model validity, we performed parameters recovery on Model 2 by simulating 130 fake participants with parameter values pseudo-randomly drawn, ensuring that parameters distributions were uncorrelated (<1%). We fitted simulated static blocks with Model 2 and compared fitted with generative parameter values. All correlations between generative and fitted parameters exceeded 63% (γ_*sen*_: 63%, σ_*inf*_: 77%, α: 88%, σ_*rep*_: 74%, γ_*rep*_ = 65%) showing a high reliability of fitted parameters estimate. Moreover, fitted parameters were uncorrelated to each other (<10%) ensuring that all parameters are correctly dissociable and do not suffer from tradeoff during the fitting procedure. **B.** To further prove model capacity in capturing participants behavior, we simulated each participant based on its own best fitting parameters. We applied to this simulated dataset the same Von Mises procedure applied to the real data and compared location (*θ*), and concentration (*κ*) parameters obtained. We found that the simulated participants faithfully replicated human data, with high correlation between simulated and real behavioral metrics. The same analysis was performed on the switch blocks with Model 4. All those controls ensure the high reliability of our models in capturing specific aspects of humans’ behavior.

**Figure S4:**
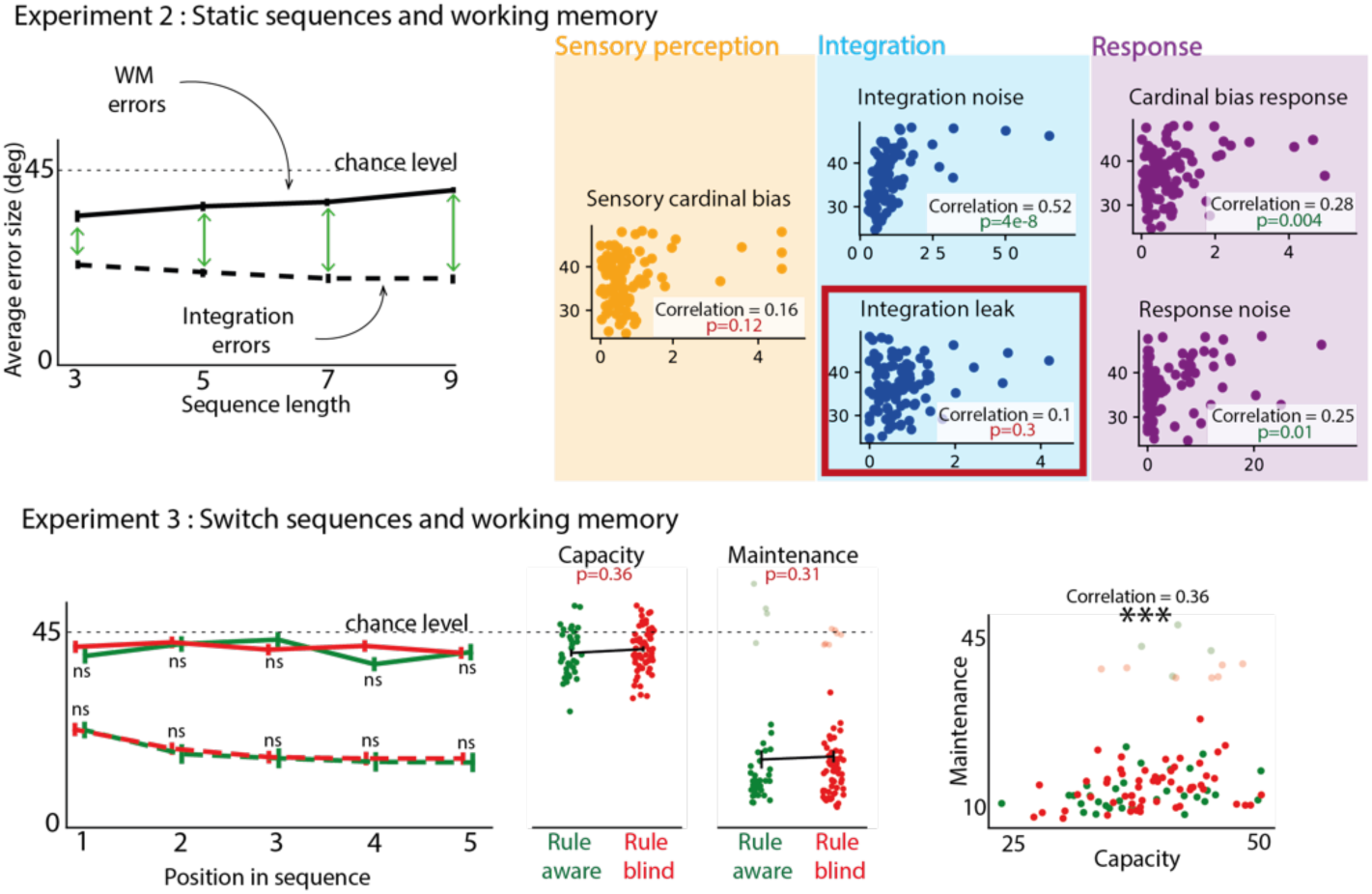
**Experiment 2**: Individual working memory performances have no impact on participants leak parameter during integration. In this experiment, 100 participants recruited online via the Prolific platform performed a task with sensory integration blocks (corresponding to the static condition of Experiment 1) and working memory blocks in which participants were presented sequences of stimuli and asked on each trial to report the orientation of one of the stimuli in the sequence (selected randomly on each trial). In both conditions, sequences could contain 3, 5, 7 or 9 oriented stimuli. Orientations were drawn uniformly in working memory blocks so that the orientation of a stimulus could not be predicted (by integration) from the orientation of another stimulus better than chance. We found that participants accuracy (average error size compared to true answer) decreased with sequence length, showing effective accumulation of evidence when presented with more information, while the same measure increased with sequence length for the working memory task, as it becomes harder to retain all elements. This suggests that the two capacities are unrelated as they qualitatively differ with sequence length. Furthermore, we fitted M2 on integration data and correlated fitted parameters with working memory capacity (average errors across all working memory trials). Importantly, we found no correlation between participants leak in integration trials and participants working memory capacity. This first additional dataset shows that sensory integration leak does not covary with working memory capacity – and likely not with general attention, since individual differences in attention should have triggered correlations in accuracy between sensory integration and working memory blocks. **Experiment 3**: To rule out the possibility that differences in working memory or general attention could explain latent rule discovery in the Switch condition. We designed a paradigm which consisted of sensory integration blocks with the same latent rule as in Experiment 1 (corresponding to the Switch condition) and two types of working memory blocks which measured participants’ ability to memorize the orientation of stimuli at positions 1-5 in the sequence (working memory capacity block) and participants’ ability to maintain the orientation of a cued stimulus at positions 1-5 in the sequence (working memory maintenance block). This test the hypothesis that participants that learn the rule might be better at remembering elements prior to the switch (elements 1-5). We collected 102 participants on this paradigm. Participants’ accuracy correlated between the two types of working memory blocks, showing accurate working memory capacity measure, but importantly did not predict latent rule discovery in sensory integration blocks. Together, those two control experiments show that differences in working memory capacities or general attention do not explain rule discovery nor leak differences between subjects.

**Figure S5:**
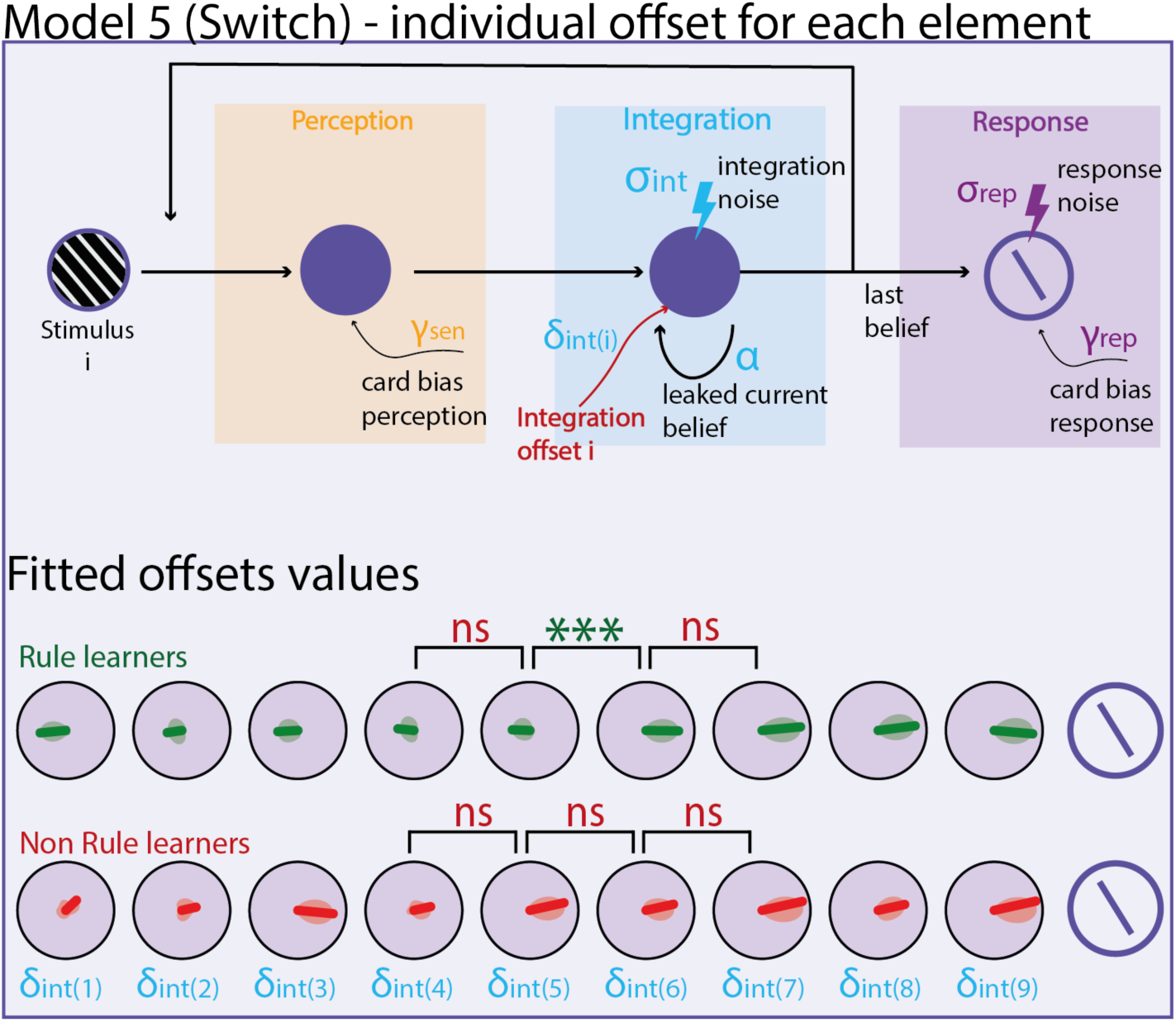
This model extends Model 4 but with an integration offset for each stimulus of the sequence. This new model confirms the validity of model 4 as all offsets of the first five stimuli are the same (∼90°) with a clear switch between stimuli 5 and 6. Participants that didn’t learn the rule, did not apply any offset to any stimulus of the sequence.

**Figure S6:**
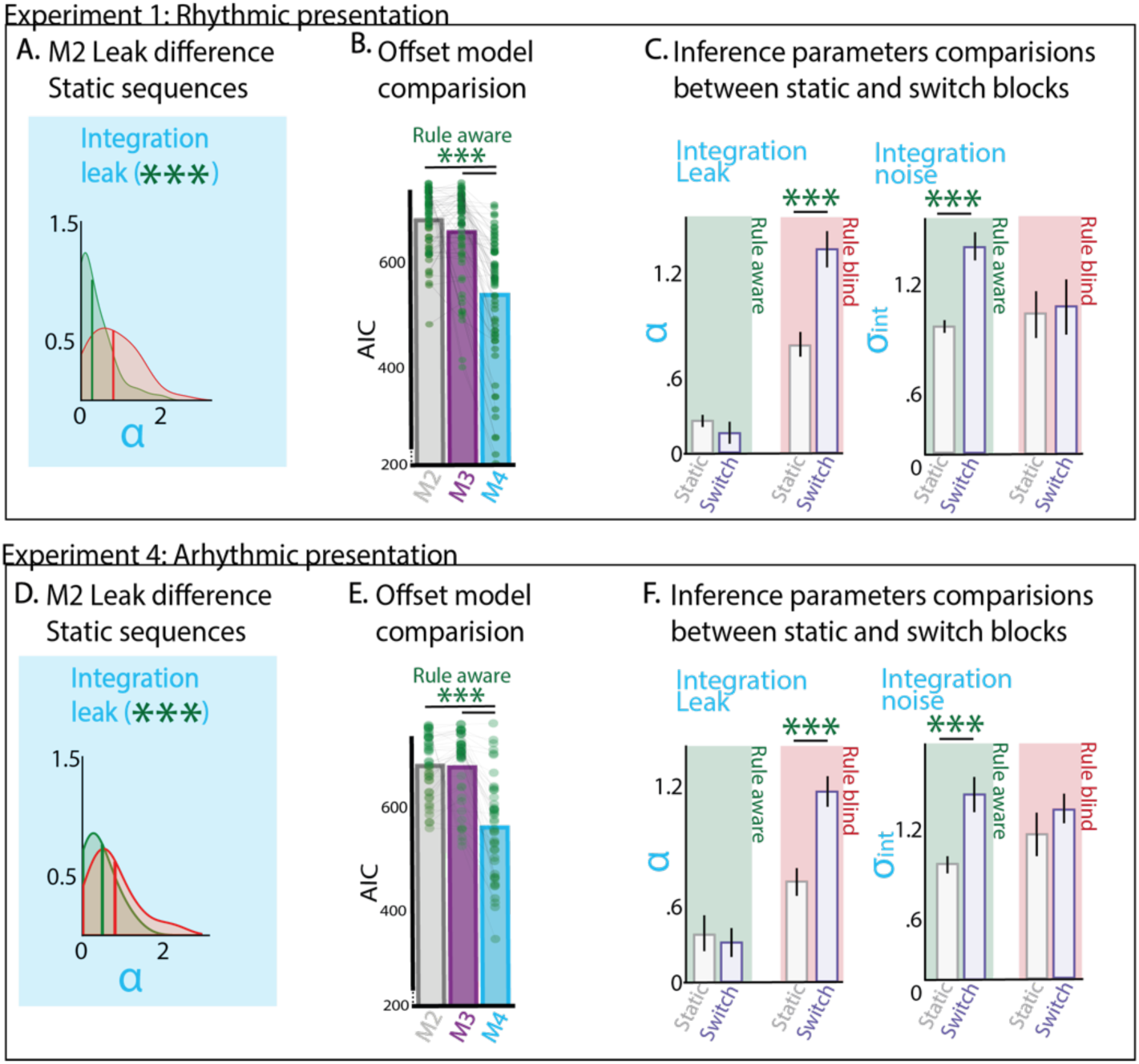
Replication of Experiment 1 (Rhythmic) main effects on a second dataset with temporal stochasticity (Experiment 4). All Main effects (Leak difference in static sequences **A&D**, offset during the integration **B&E** and modification of integration parameters when uncovering the hidden rule **C&F**) have been replicated in this second dataset.

**Figure S7:**
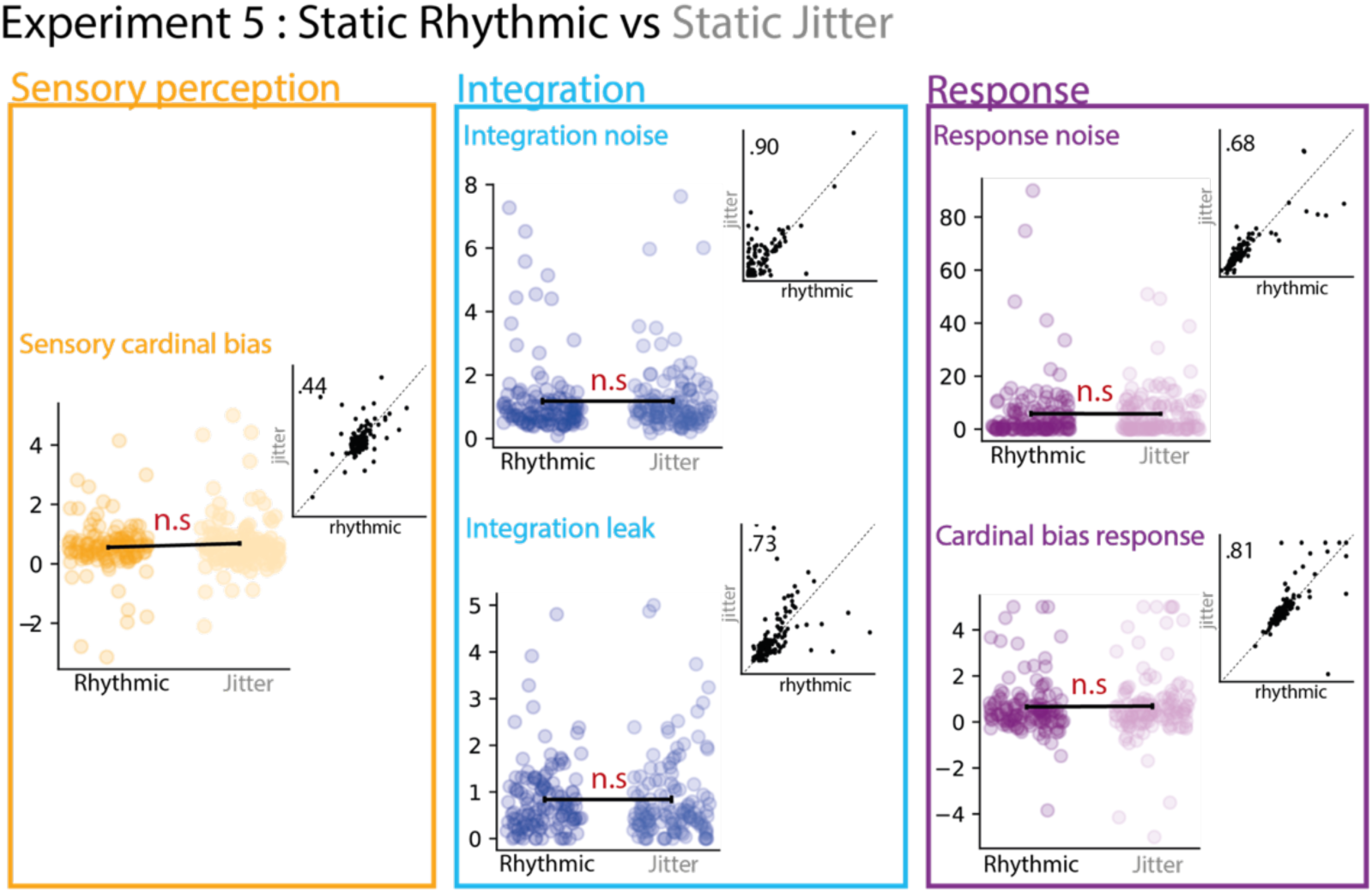
In Experiment 5, 126 participants performed a task only composed of static sequences but in blocks of rhythmic or jittered presentation. Fitting Model 2 on data from rhythmic or jittered trials only yield no differences. Moreover, we found important correlation between individual parameters fitted for rhythmic and for jittered data showing a good test re-test reliability of these parameters despite the different temporal predictability of elements. This experiment confirms the absence of impact of temporal stochasticity on integration parameters already suggested by the comparison of static sequences from Experiment 1 & and 4 (Fig 4A).

**Figure S8:**
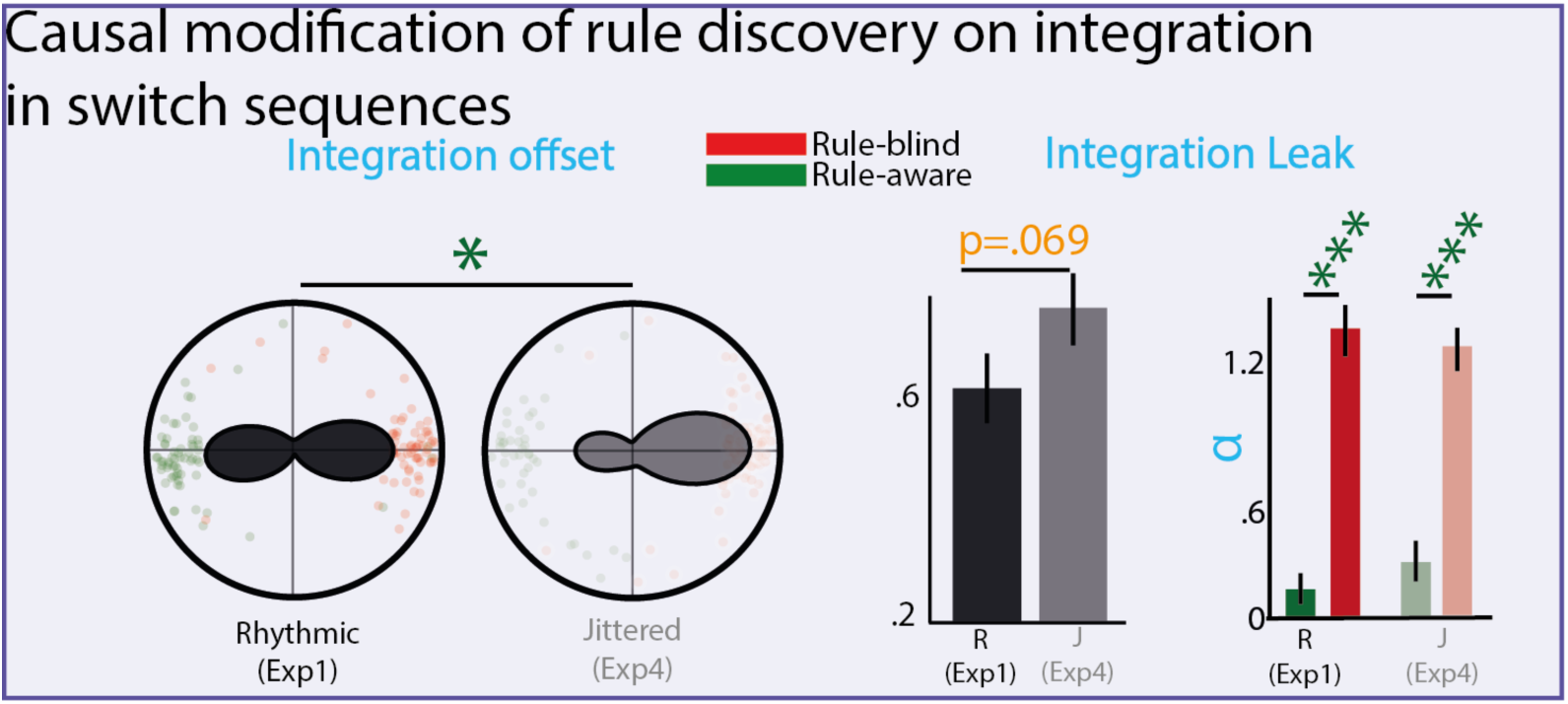
Comparison of jittered vs rhythmic integration parameters on switch sequences. **Left**: Integration offset difference between rhythmic and jittered experiments **Right**: Integration leak difference between rhythmic and jittered experiments. Informed by the analysis of late rule-aware subjects in Experiment 1, we reasoned that if rule discovery causally influences integration during switch sequences, then: 1/ the distribution of integration offsets should differ between the two experiments, and 2/ the integration leak should be larger during switch blocks in the jittered (exp 4) compared to the rhythmic experiment (Exp 1). Post-hoc tests confirmed these hypotheses for the offset distributions (p<0.05), and the leak (marginal significance: one sided p=0.069-), supporting findings from late rule-aware subjects. In both Experiments 1 and 4, the leak was very significantly different between rule-aware and rule-blind subjects (both ps<0.001), and similar across experiments, suggesting that the rhythmic vs jittered leak difference is solely driven by the smaller number of rule-aware participants in the jittered experiment.

**Figure S9:**
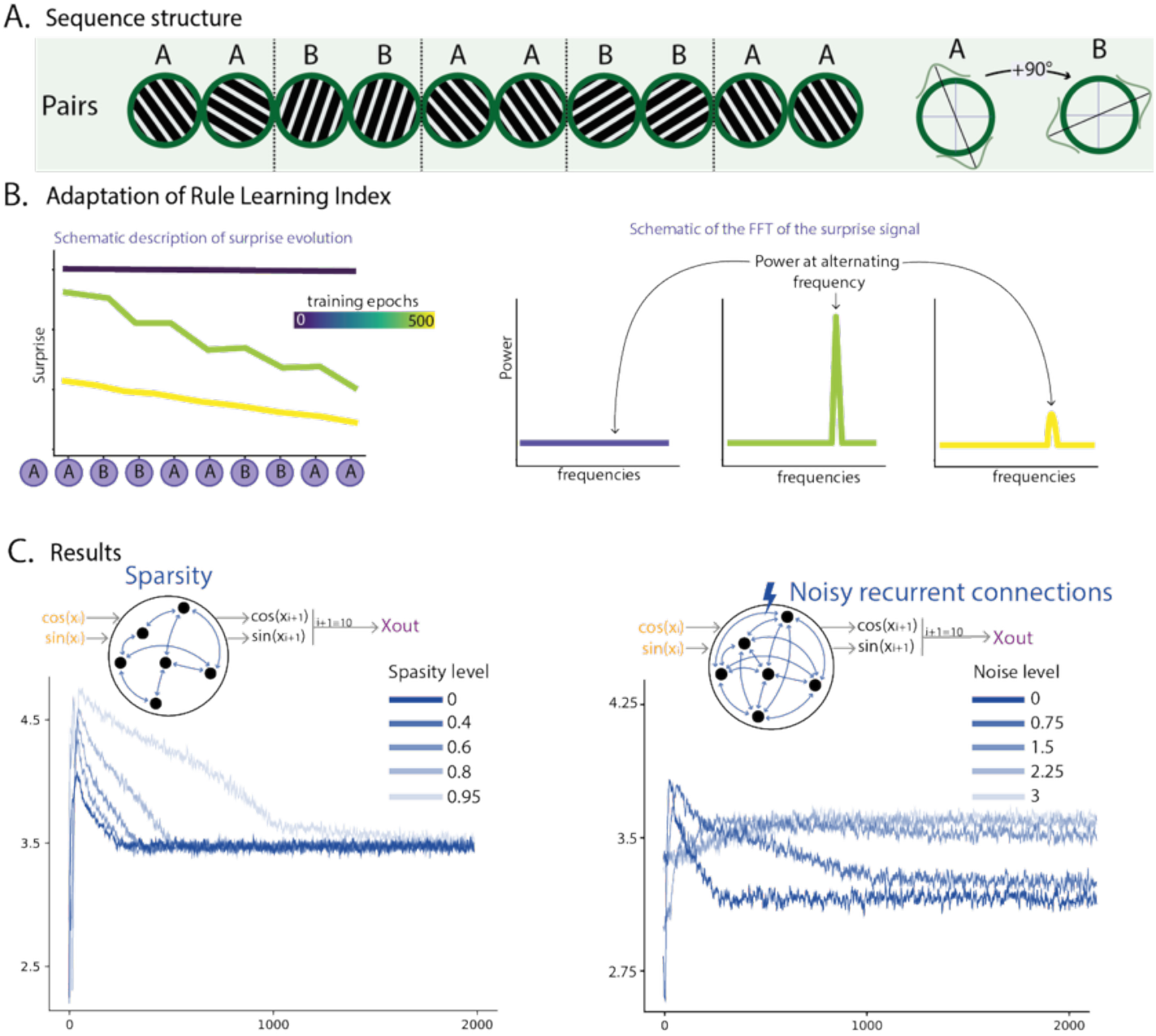
replication of the effect of sparsity and noise in recurrent connection on the effect of rule discovery with a different rule sequence. **A.** Description of the AABBAABBAA sequence. **B.** We adapted the rule learning index defined in the main text to measure the learning of the pairs. When the rule is not yet grasped by the network, the surprise vector shows a periodic pattern at the period of the pairs. We computed the FFT of the surprise signal and retrieved the power at the frequency of the pair. The evolution of the power at this specific frequency reflects how much the network is surprised at each switch. **C.** We found qualitatively similar results to what was found for the switch rule: the sparsity of the networks influenced the time needed for the rule discovery while the noisy recurrent connections influences both the integration and the rule learning.

**Figure S10:**
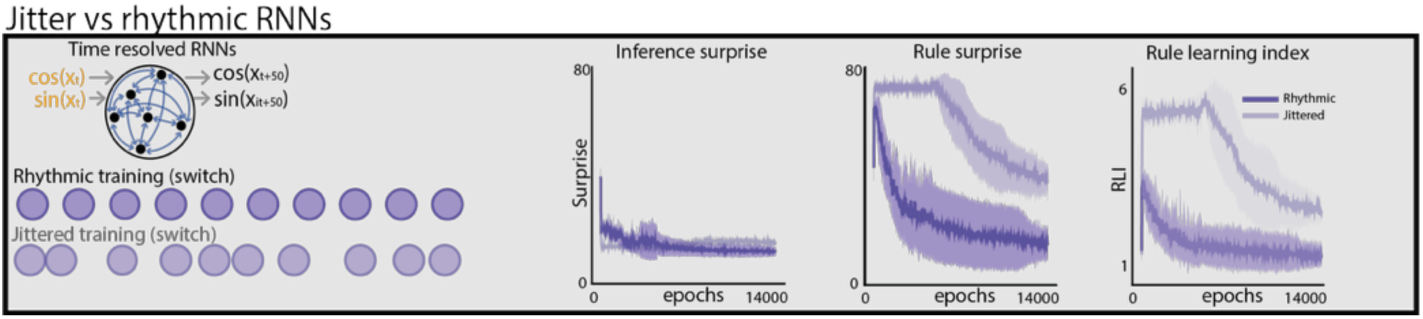
Details on the analysis of time-resolved RNNs. To adapt our RNNs to the analysis of rhythmic vs jittered sequences, we turn them to time resolved RNNs. Instead of predicting the next element, they should predict the content of the next 50ms and thus have to make correct assumptions about what (the content) and when (the correct 50ms window) to accurately predict the sequence. The loss was computed as the MSE of the actual positions of the elements in the sequence. We trained 20 RNNs of Rhythmic sequences and 20 more of jittered sequences. For each of those RNNs we recorded during Inference Surprise and Rule Surprise along the 15000 epochs of the training. Inference surprise (ie. average of the prediction errors for elements 2,3,4,5& 7,8,9,10 – like for non-time resolved RNNs) is a proxy of inference learning and plotted on the left panel. We can see that rhythmic vs jitter training marginally changes this metric: the initial learning speed is slightly increased in presence of jitter in the training material and later on (around 6000 epochs) when the networks start to discover the rule, the inference is slightly impaired revealing a tradeoff in RNNs between the two forms of inference. At the opposite, le rule surprise (prediction error consecutive to the switch, at element 6) is greatly impaired by jitter in the input. While RNNs trained with rhythmic sequences immediately start learning the rule, those trained with jittered sequences only starts learning the rule later on and slower. The last panel represents the Rule learning index = the ratio between Rule surprise and Inference Surprise.

**Figure S11:**
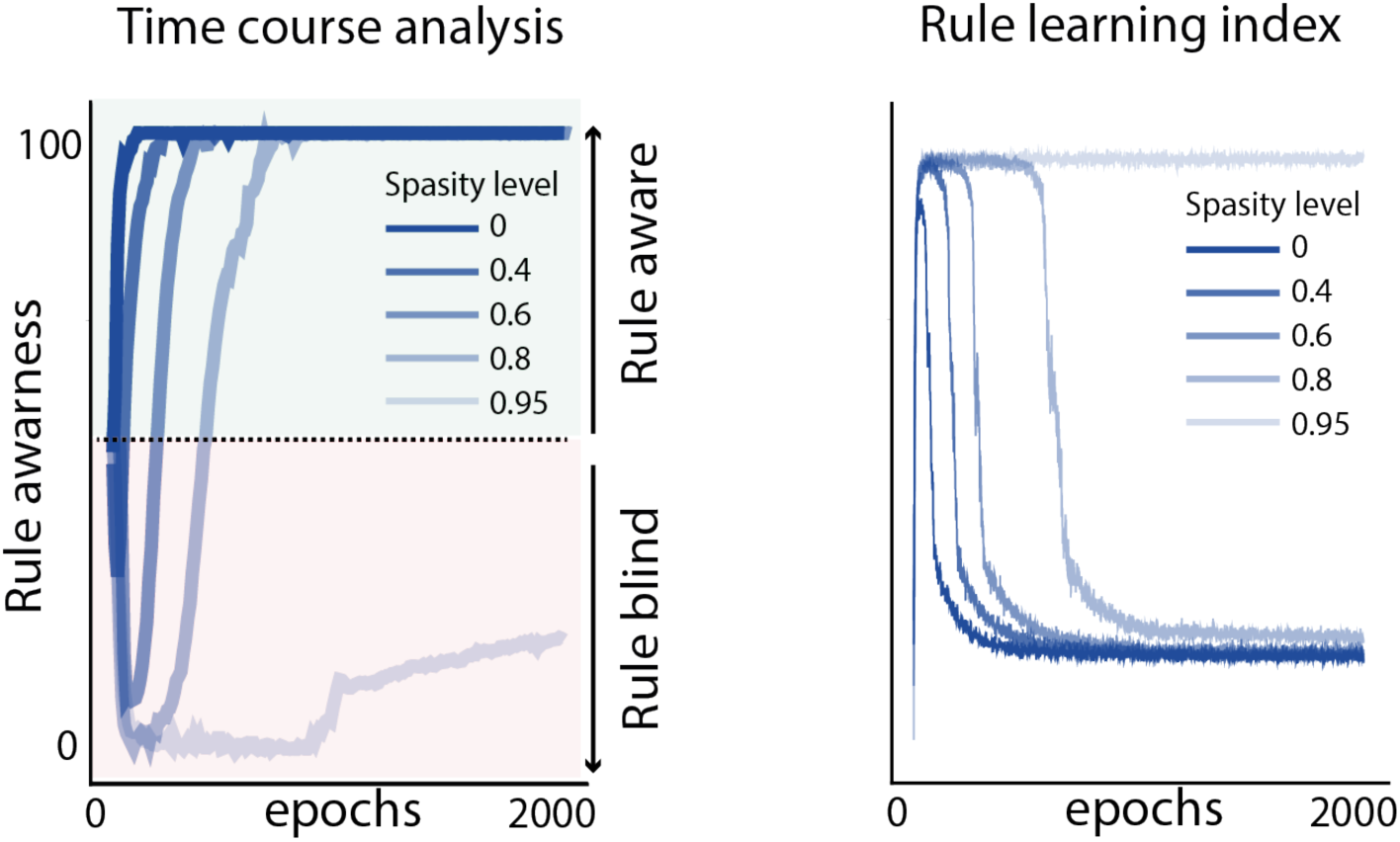
Replication of the effect of RNN sparsity on rule discovery with time course analysis. This second metric was used to better match humans and RNNs analyses. We re-applied the same procedure as for the time course of rule discovery in humans. For that, after each training epoch, we estimated the percentage of answers from t+7 and t+5 trials that are compatible with the knowledge of the rule. This gives the time course of rule discovery in RNNs similar to what was made for humans (**Fig 3C**). We replicated our main effect with this metric by varying the sparsity of the network (0, 0.4, 0.6, 0.8 & 0.95) and found that rule discovery in RNN was delayed because of the sparsity. This metric closely matches the time course analysis in humans and replicates our main effect found with RLI with a delayed rule discovery caused by higher sparsity levels in the networks.

## References

1. Bogacz, R. Optimal decision-making theories: linking neurobiology with behaviour. Trends in Cognitive Sciences 11, 118–125 (2007).

2. Morillon, B., Schroeder, C. E. & Wyart, V. Motor contributions to the temporal precision of auditory attention. Nat Commun 5, 5255 (2014).

3. Summerfield, C. & Egner, T. Expectation (and attention) in visual cognition. Trends in Cognitive Sciences 13, 403–409 (2009).

4. Wald, A. & Wolfowitz, J. Optimum Character of the Sequential Probability Ratio Test. The Annals of Mathematical Statistics 19, 326–339 (1948).

5. Janata, P., Tomic, S. T. & Haberman, J. M. Sensorimotor coupling in music and the psychology of the groove. J Exp Psychol Gen 141, 54– 75 (2012).

6. Nobre, A. C. & van Ede, F. Anticipated moments: temporal structure in attention. Nat Rev Neurosci 19, 34–48 (2018).

7. Wollman, I. & Morillon, B. Organizational principles of multidimensional predictions in human auditory attention. Sci Rep 8, 13466 (2018).

8. Zalta, A., Petkoski, S. & Morillon, B. Natural rhythms of periodic temporal attention. Nat Commun 11, 1051 (2020).

9. Cheadle, S. et al. Adaptive Gain Control during Human Perceptual Choice. Neuron 81, 1429–1441 (2014).

10. Freedman, D. J. & Assad, J. A. Neuronal Mechanisms of Visual Categorization: An Abstract View on Decision Making. Annu. Rev. Neurosci. 39, 129–147 (2016).

11. Wyart, V., de Gardelle, V., Scholl, J. & Summerfield, C. Rhythmic Fluctuations in Evidence Accumulation during Decision Making in the Human Brain. Neuron 76, 847–858 (2012).

12. Benjamin, L., Sablé-Meyer, M., Fló, A., Dehaene-Lambertz, G. & Roumi, F. A. Long-Horizon Associative Learning Explains Human Sensitivity to Statistical and Network Structures in Auditory Sequences. J. Neurosci. 44, e1369232024 (2024).

13. Benjamin, L., Fló, A., Al Roumi, F. & Dehaene-Lambertz, G. Humans parsimoniously represent auditory sequences by pruning and completing the underlying network structure. eLife 12, e86430 (2023).

14. Benjamin, L. et al. Tracking transitional probabilities and segmenting auditory sequences are dissociable processes in adults and neonates. Developmental Science 26, e13300 (2023).

15. Henin, S. et al. Learning hierarchical sequence representations across human cortex and hippocampus. Science Advances 7, 1–13 (2021).

16. Saffran, J. R., Aslin, R. N. & Newport, E. L. Statistical Learning by 8-Month-Old Infants. Science 274, 1926–1928 (1996).

17. Schapiro, A. C., Turk-Browne, N. B., Norman, K. A. & Botvinick, M. M. Statistical learning of temporal community structure in the hippocampus. Hippocampus 26, 3–8 (2016).

18. Schapiro, A. C., Rogers, T. T., Cordova, N. I., Turk-, N. B. & Botvinick, M. M. Neural representations of events arise from temporal community structure. Nature Neuroscience 16, 486–492 (2013).

19. Al Roumi, F., Marti, S., Wang, L., Amalric, M. & Dehaene, S. Mental compression of spatial sequences in human working memory using numerical and geometrical primitives. Neuron 109, 2627–2639.e4 (2021).

20. Al Roumi, F., Planton, S., Wang, L. & Dehaene, S. Brain-imaging evidence for compression of binary sound sequences in human memory. eLife 12, e84376 (2023).

21. Albouy, P., Martinez-Moreno, Z. E., Hoyer, R. S., Zatorre, R. J. & Baillet, S. Supramodality of neural entrainment: Rhythmic visual stimulation causally enhances auditory working memory performance. Science Advances 8, eabj9782 (2022).

22. Quilty-Dunn, J., Porot, N. & Mandelbaum, E. The Best Game in Town: The Re-Emergence of the Language of Thought Hypothesis Across the Cognitive Sciences. Behavioral and Brain Sciences 1–55 (2022) doi:10.1017/S0140525X22002849.

23. Xie, Y. et al. Geometry of sequence working memory in macaque prefrontal cortex. Science 375, 632–639 (2022).

24. Fiser, J. & Lengyel, G. Statistical Learning in Vision. Annu. Rev. Vis. Sci. 8, 265–290 (2022).

25. Ma, W. J., Kording, K. & Goldreich, D. Bayesian Models of Perception and Action: An Introduction. (The MIT Press, London, England, 2023).

26. Ratcliff, R. & McKoon, G. The Diffusion Decision Model: Theory and Data for Two-Choice Decision Tasks. Neural Comput 20, 873–922 (2008).

27. Beck, J. M., Ma, W. J., Pitkow, X., Latham, P. E. & Pouget, A. Not Noisy, Just Wrong: The Role of Suboptimal Inference in Behavioral Variability. Neuron 74, 30–39 (2012).

28. Drugowitsch, J., Wyart, V., Devauchelle, A.-D. & Koechlin, E. Computational Precision of Mental Inference as Critical Source of Human Choice Suboptimality. Neuron 92, 1398–1411 (2016).

29. Findling, C. & Wyart, V. Computation noise in human learning and decision-making: origin, impact, function. Current Opinion in Behavioral Sciences 38, 124–132 (2021).

30. Rahnev, D. & Denison, R. N. Suboptimality in perceptual decision making. Behav Brain Sci 41, e223 (2018).

31. Dehaene, S., Al Roumi, F., Lakretz, Y., Planton, S. & Sablé-Meyer, M. Symbols and mental programs: a hypothesis about human singularity. Trends in Cognitive Sciences 26, 751–766 (2022).

32. Wu, S., Thalmann, M. & Schulz, E. Two types of motifs enhance human recall and generalization of long sequences. Commun Psychol 3, 3 (2025).

33. Kleinman, M., Chandrasekaran, C. & Kao, J. A mechanistic multi-area recurrent network model of decision-making. in Advances in Neural Information Processing Systems vol. 34 23152–23165 (Curran Associates, Inc., 2021).

34. Kurikawa, T. Different timescales of neural activities introduce different representations of task-relevant information. Preprint at 10.1101/2024.07.23.604720 (2024).

35. Lin, B., Bouneffouf, D. & Cecchi, G. Predicting human decision making in psychological tasks with recurrent neural networks. PLoS ONE 17, e0267907 (2022).

36. Girshick, A. R., Landy, M. S. & Simoncelli, E. P. Cardinal rules: visual orientation perception reflects knowledge of environmental statistics. Nat Neurosci 14, 926–932 (2011).

37. Lee, J. K., Rouault, M. & Wyart, V. Adaptive tuning of human learning and choice variability to unexpected uncertainty. SCIENCE ADVANCES.

38. Ossmy, O. et al. The timescale of perceptual evidence integration can be adapted to the environment. Curr Biol 23, 981–986 (2013).

39. Gómez Varela, I., Orpella, J., Poeppel, D., Ripolles, P. & Assaneo, M. F. Syllabic rhythm and prior linguistic knowledge interact with individual differences to modulate phonological statistical learning. Cognition 245, 105737 (2024).

40. Maheu, M., Meyniel, F. & Dehaene, S. Rational arbitration between statistics and rules in human sequence learning. 10.1101/2020.02.06.937706 (2020) doi:10.1101/2020.02.06.937706.

41. Waskom, M. L. & Kiani, R. Decision Making through Integration of Sensory Evidence at Prolonged Timescales. Current Biology 28, 3850–3856.e9 (2018).

42. Wyart, V. & Koechlin, E. Choice variability and suboptimality in uncertain environments. Current Opinion in Behavioral Sciences 11, 109–115 (2016).

43. Eckstein, M. K., Summerfield, C., Daw, N. D. & Miller, K. J. Predictive and Interpretable: Combining Artificial Neural Networks and Classic Cognitive Models to Understand Human Learning and Decision Making. 2023.05.17.541226 Preprint at 10.1101/2023.05.17.541226 (2023).

44. Peterson, J. C., Bourgin, D. D., Agrawal, M., Reichman, D. & Griffiths, T. L. Using large-scale experiments and machine learning to discover theories of human decision-making. Science 372, 1209–1214 (2021).

45. Findling, C. & Wyart, V. Computation noise promotes zero-shot adaptation to uncertainty during decision-making in artificial neural networks. Science Advances 10, eadl3931 (2024).

46. Flesch, T., Juechems, K., Dumbalska, T., Saxe, A. & Summerfield, C. Orthogonal representations for robust context-dependent task performance in brains and neural networks. Neuron 110, 1258–1270.e11 (2022).

47. Molano-Mazón, M. et al. Recurrent networks endowed with structural priors explain suboptimal animal behavior. Curr Biol 33, 622–638.e7 (2023).

48. Deneve, S., Latham, P. E. & Pouget, A. Efficient computation and cue integration with noisy population codes. Nat Neurosci 4, 826– 831 (2001).

49. Ermentrout, B. Neural networks as spatio-temporal pattern-forming systems. Rep. Prog. Phys. 61, 353 (1998).

50. Esnaola-Acebes, J. M., Roxin, A. & Wimmer, K. Bump attractor dynamics underlying stimulus integration in perceptual estimation tasks. 2021.03.15.434192 Preprint at 10.1101/2021.03.15.434192 (2021).

51. Jazayeri, M. & Movshon, J. A. Optimal representation of sensory information by neural populations. Nat Neurosci 9, 690–696 (2006).

52. Pouget, A., Dayan, P. & Zemel, R. S. Inference and computation with population codes. Annual Review of Neuroscience 26, 381–410 (2003).

53. Garvert, M. M., Dolan, R. J. & Behrens, T. E. J. A map of abstract relational knowledge in the human hippocampal–entorhinal cortex. eLife 6, 1–20 (2017).

54. Planton, S. et al. A theory of memory for binary sequences: Evidence for a mental compression algorithm in humans. PLoS Computational Biology 10.1371/journal.pcbi.1008598 (2021) doi:10.1371/journal.pcbi.1008598.

55. Wyart, V., Myers, N. E. & Summerfield, C. Neural Mechanisms of Human Perceptual Choice Under Focused and Divided Attention. J. Neurosci. 35, 3485–3498 (2015).

56. Schuck, N. W. et al. Medial Prefrontal Cortex Predicts Internally Driven Strategy Shifts. Neuron 86, 331–340 (2015).

57. Hamilton, D. A. & Brigman, J. L. Behavioral flexibility in rats and mice: contributions of distinct frontocortical regions. Genes Brain and Behavior 14, 4–21 (2015).

58. Costa, V. D., Tran, V. L., Turchi, J. & Averbeck, B. B. Reversal Learning and Dopamine: A Bayesian Perspective. J. Neurosci. 35, 2407– 2416 (2015).

59. Clark, L., Cools, R. & Robbins, T. W. The neuropsychology of ventral prefrontal cortex: Decision-making and reversal learning. Brain and Cognition 55, 41–53 (2004).

60. Glaze, C. M., Kable, J. W. & Gold, J. I. Normative evidence accumulation in unpredictable environments. eLife 4, e08825 (2015).

61. Izquierdo, A., Brigman, J. L., Radke, A. K., Rudebeck, P. H. & Holmes, A. The neural basis of reversal learning: An updated perspective. Neuroscience 345, 12–26 (2017).

62. Mante, V., Sussillo, D., Shenoy, K. V. & Newsome, W. T. Context-dependent computation by recurrent dynamics in prefrontal cortex. Nature 503, 78–84 (2013).

63. Pestilli, F., Carrasco, M., Heeger, D. J. & Gardner, J. L. Attentional Enhancement via Selection and Pooling of Early Sensory Responses in Human Visual Cortex. Neuron 72, 832–846 (2011).

64. Dubreuil, A., Valente, A., Beiran, M., Mastrogiuseppe, F. & Ostojic, S. The role of population structure in computations through neural dynamics. Nat Neurosci 25, 783–794 (2022).

65. Morillon, B. & Baillet, S. Motor origin of temporal predictions in auditory attention. Proceedings of the National Academy of Sciences of the United States of America 114, E8913–E8921 (2017).

66. Waskom, M. L. & Kiani, R. Decision Making through Integration of Sensory Evidence at Prolonged Timescales. Current Biology 28, 3850–3856.e9 (2018).

67. de Leeuw, J. R. jsPsych: A JavaScript library for creating behavioral experiments in a Web browser. Behav Res 47, 1–12 (2015).

68. Usher, M. & McClelland, J. L. The time course of perceptual choice: the leaky, competing accumulator model. Psychol Rev 108, 550–592 (2001).

69. Drevet, J., Drugowitsch, J. & Wyart, V. Efficient stabilization of imprecise statistical inference through conditional belief updating. Nat Hum Behav 6, 1691–1704 (2022).

70. Lee, J. K., Rouault, M. & Wyart, V. Adaptive tuning of human learning and choice variability to unexpected uncertainty. SCIENCE ADVANCES.

71. Acerbi, L. & Ma, W. J. Practical Bayesian Optimization for Model Fitting with Bayesian Adaptive Direct Search. Preprint at 10.48550/arXiv.1705.04405 (2017).

